# Immunoinformatic and dynamic simulation-based designing of a multi-epitope vaccine against emerging pathogen *Elizabethkingia anophelis*

**DOI:** 10.1101/758219

**Authors:** Zulkar Nain, Faruq Abdullah, M. Mizanur Rahman, Mohammad Minnatul Karim, S. M. Raihan Rahman, Shakil Ahmed Khan, Sifat Bin Sayed, Md. Moinuddin Sheam, Zahurul Haque, Utpal Kumar Adhikari

## Abstract

*Elizabethkingia anophelis* is an emerging human pathogen causing neonatal meningitis, catheter-associated infections and nosocomial outbreaks with high mortality rates. Besides, they are resistant to most antibiotics used in empirical therapy. In this study, therefore, we used immunoinformatic approaches to design an epitope-based vaccine against *E. anophelis* as an alternative preventive measure. Initially, T-cell (CTL and HTL) and B-cell (LBL) epitopes were predicted from the highest antigenic protein. The CTL and HTL epitopes together had a population coverage of 99.97% around the world. Eventually, 6 CTL, 7 HTL, and 2 LBL epitopes were selected and used to construct a multiepitope vaccine. The vaccine protein was found to be highly immunogenic, non-allergenic, and non-toxic. Codon adaptation and *in silico* cloning were performed to ensure better expression within *E. coli* K12 host system. The stability of the vaccine structure was also improved by disulphide bridging. In addition, molecular docking and dynamic simulation revealed good and stable binding affinity between the vaccine and receptor. The immune simulation showed higher levels of T-cell and B-cell activities which was in coherence with actual immune response. Repeated exposure simulation resulted in higher clonal selection and faster antigen clearance. Nevertheless, experimental validation is required to ensure the immunogenic potency and safety of this vaccine to control *E. anophelis* infection in the future.

## Background

*Elizabethkingia anophelis* is an opportunistic human pathogen responsible for neonatal meningitis^1,2^ and infections in immunocompromised^3,4^ and postoperative patients^5^. These are Gram-negative bacilli belonging to the family of *Flavobacteriaceae*^1,6^. Despite their widespread presence^1^ and rare association with human infection, it has recently been reported that bacteria of the genus *Elizabethkingia* are involved in life-threatening infections^7^. The first isolation of *E. anophelis* from the midgut of mosquito *Anophelis gambiae* was dated back to 2011^8^. Over the last five years, *E. anophelis* has caused multiple outbreaks in South Africa^9^, Singapore^10^, Hong Kong^11^, Taiwan^7^ and the United States^12^ with a mortality of up to 60%^7,11,13^. For instance, two large outbreaks in the Midwestern United States resulted in a death toll of 31% (20/65) in 65 cases^13,14^. In addition, in Illinois, at least 10 cases of *E. anophelis* infections were reported from January 2014 to April 2016 with six (60%) of the patients dying from this infection^3^. The significant mortality outcome from *E. anophelis* infections may be correlated with the virulence factors (*i*.*e*., capsule, lipopolysaccharide, magnesium transport protein, etc.) of the bacteria and pre-existing health conditions of the sufferers^7^. Also, multiple antimicrobial resistance is an integral part of the *E. anophelis* genome, rendering it immune to the last line of antibiotics used in empirical therapy^1^.

Under the genus *Elizabethkingia, E. meningoseptica* is well-documented as a deadly infectious agent, which can also cause meningitis and sepsis as well as nosocomial outbreaks with a mortality rate of about 50%^5^. Interestingly, recent studies indicated that many of the previously described *E. meningoseptica* were indeed *E. anophelis*^7^. Therefore, a significant portion of the previous *Elizabethkingia* infections could actually be *E. anophelis* infections^11^. As a result, *E. anophelis* has been considered as an emerging opportunistic pathogen whose clinical significance has recently been emphasized^1^. However, little is known about this newly emerging pathogen. For instance, the transmission route of *E. anophelis* still remains elusive^6,15^. These altogether have made conventional treatment against *E. anophelis* infections obsolete and challenging^2^. Nevertheless, there are no vaccines or treatments available to prevent and/or control the diseases associated with *E. anophelis*. Therefore, alternative control measures are essential to counteract their pathogenic potential.

Peptide vaccines are a newer form of immune stimulant where fragments (*i*.*e*., epitopes) of antigenic protein are used to mimic the natural presence of pathogens^16^. In 1985, the first epitope-based vaccine was developed against cholera in *Escherichia coli*^17^. Besides, there are many peptide vaccines under development, such as the vaccine for HIV, malaria, swine fever, influenza, anthrax, etc.^16^. The developmental process includes the identification of the virulence protein, selection of peptide segments and generation of cellular and humoral immune response^18^. The protein antigen in *E. anophelis* proteome can be identified from the recently sequenced *E. anophelis* Ag1 genome^19^. Interestingly, computer-based advanced approaches like immunoinformatics can be useful in this scenario. In recent years, the use of immunoinformatics to design *in silico* peptide vaccine model against viruses^20–26^ bacteria^27–33^ and parasites^34–36^ have been intensified. Therefore, peptide vaccine can be a more attractive alternative as preventive means against *E. anophelis*.

The current study involves the screening of the *E. anophelis* proteome to identify the highest antigenic protein, followed by the prediction of different T and B-cell epitopes with their corresponding MHC alleles. These epitopes were further evaluated prior to estimating population coverage. Therefore, a multiepitope vaccine was designed using the most potent epitopes with appropriate adjuvant and linkers. The primary sequence was used for the immunogenic and physiochemical profiling of the vaccine protein and for the prediction of secondary and tertiary structure. The predicted 3D structure was then refined and subjected to disulphide engineering to improve stability. The binding interaction of the vaccine to the receptor was calculated by molecular docking. Finally, molecular dynamics and immune simulation were performed to estimate the stability of the vaccine and its immunogenic potency in real-life, respectively. The overall computational workflow of the study is graphically illustrated in Fig. 1.

**Figure 1:**
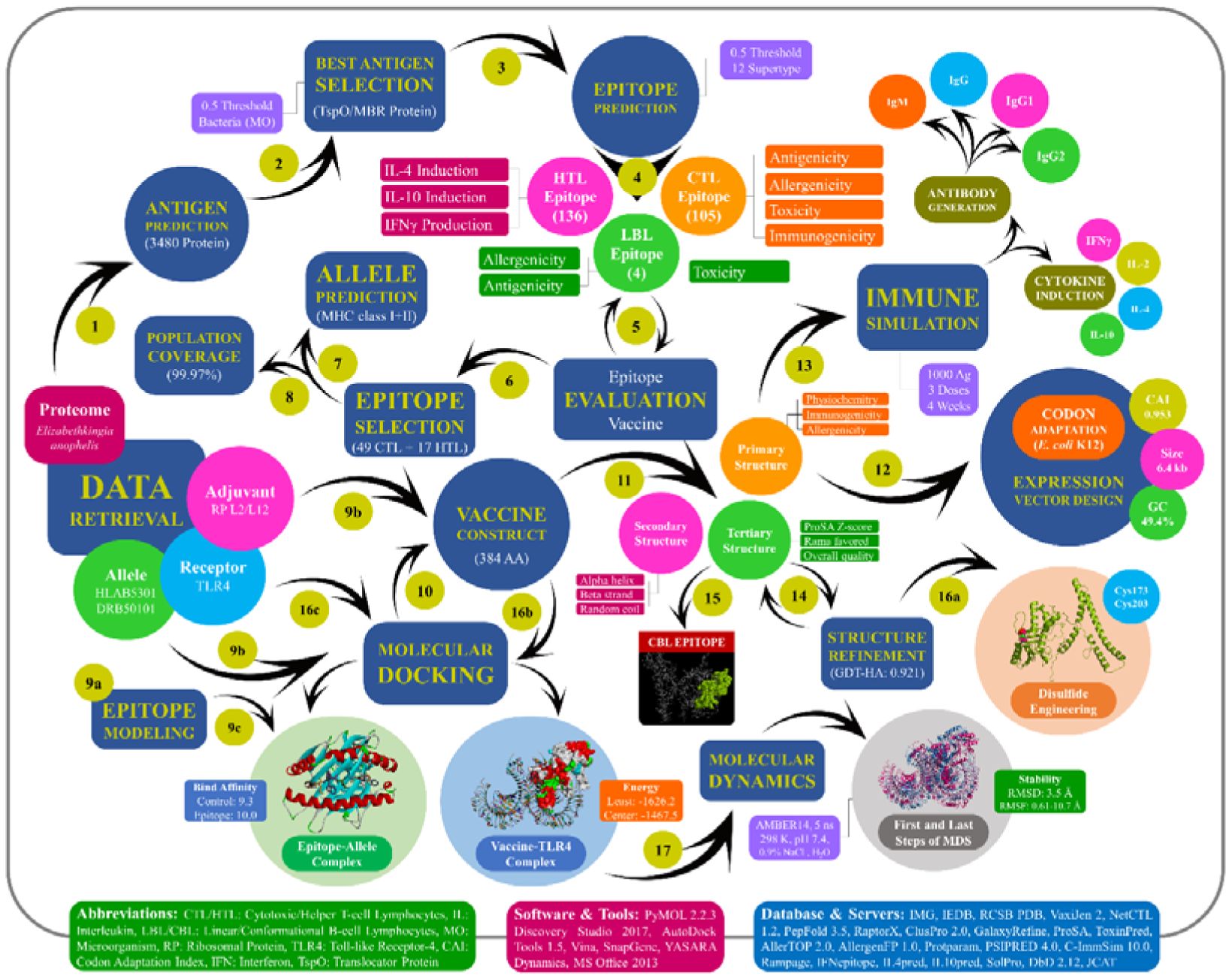
Experimental workflow used to develop a multiepitope vaccine against emerging bacterial pathogen *E. anophelis*. Here, boxes represent the methods and parameters while the arrow number shows the study steps in sequence.

## Results

### Highest antigenic protein selection

Out of total 3648 proteins in *E. anophelis* Ag1 proteome, 3480 proteins were left after subtracting genes with homologs in *Homo sapiens* in a total 179 hits in the IMG/M database. The VaxiJen 2.0 server predicts antigenic potency by assigning each protein a score. The higher the score, the higher the antigenicity. The highest score (1.0568) was found for TSPO/MBR related protein (GenBank ID: WP_009087390) and therefore selected for the design of a multiepitope vaccine. This protein is a translocator protein (TSPO), also known as tryptophan-rich sensory protein, which contains 162 amino acid residues.

### Prediction and assessment of CTL, HTL and LBL epitopes

A total of 105 unique CTL epitopes (9-mer) were predicted from the *E. anophelis* TSPO protein using the NetCTL 1.2 server. Among them, only 49 epitopes were found as antigenic, immunogenic, non-allergenic and non-toxic as tabulated in Supplementary Table 1. These epitopes were further used to predict their MHC-I binding alleles using MHC-I allele prediction tool of the IEDB server. Similarly, a total of 136 unique HTL epitopes (15-mer) and their MHC-II binding molecules were predicted using the IEDB MHC-II prediction tool. Further, the cytokine (*i*.*e*., IFN-γ, IL-4, and IL-10) inducing ability of these HTL epitopes were evaluated. However, only 17 HTL epitopes were able to induce all three types of cytokines as shown in Supplementary Table 2. B-cell epitopes are antigenic regions of a protein that can trigger antibody formation. We found 4 linear B lymphocyte (LBL) epitopes in of variable length ranged in 8 to 26 amino acid residues while their scores ranged from 0.686 to 0.794. All predicted LBL epitopes, along with their length and scores, are provided in Supplementary >Table 3. After evaluation, however, only 2 LBL epitopes (*i*.*e*., MKRQIDWKALAA and ALTPANSFNVWYNQLDKPLFTPPSNF) were found to be antigenic, non-allergenic and non-toxic, hence, considered for vaccine construction.

### Population coverage analysis

The finally selected 49 CTL and 17 HTL epitopes and their corresponding HLA alleles were used to estimate the population coverage both individually and in combination as tabulated in Supplementary Table 4. Our selected CTL and HTL epitopes were found to cover 99.92% and 68.89% of the world population, respectively. In this study, we attached importance to the combined population coverage, as both CTL and HTL epitopes were considered in the vaccine to construct. Importantly, CTL and HTL epitopes showed 99.97% population coverage when used in combination. The highest population coverage (100%) was found for several European countries (*viz*., Bulgaria, England, Finland, France, Germany, Poland, and Sweden) and the South American country Peru. The least population coverage (0%) was predicted for the European country Slovakia, which is plausible due to its lower allele frequency (based on the data available in Allele Frequencies (http://allelefrequencies.net) database). In West Africa, where the bacteria first appeared, the population coverage was 99.81%. *E. anophelis* caused several outbreaks in different countries of the world, especially in the USA. Therefore, population coverage in these regions is important for vaccine development. Importantly, the cumulative population coverages found in the outbreak region USA, South Africa, China, Singapore, Hong Kong, and Taiwan were 99.98%, 99.51%, 99.39%, 99.44%, 97.96%, and 99.71%, respectively as depicted in Fig. 2. The result suggested that our vaccine could help to control *E. anophelis* in these regions. In addition, the population coverage in South Asia and Australia were found to be 99.86% and 99.30%, respectively.

**Figure 2:**
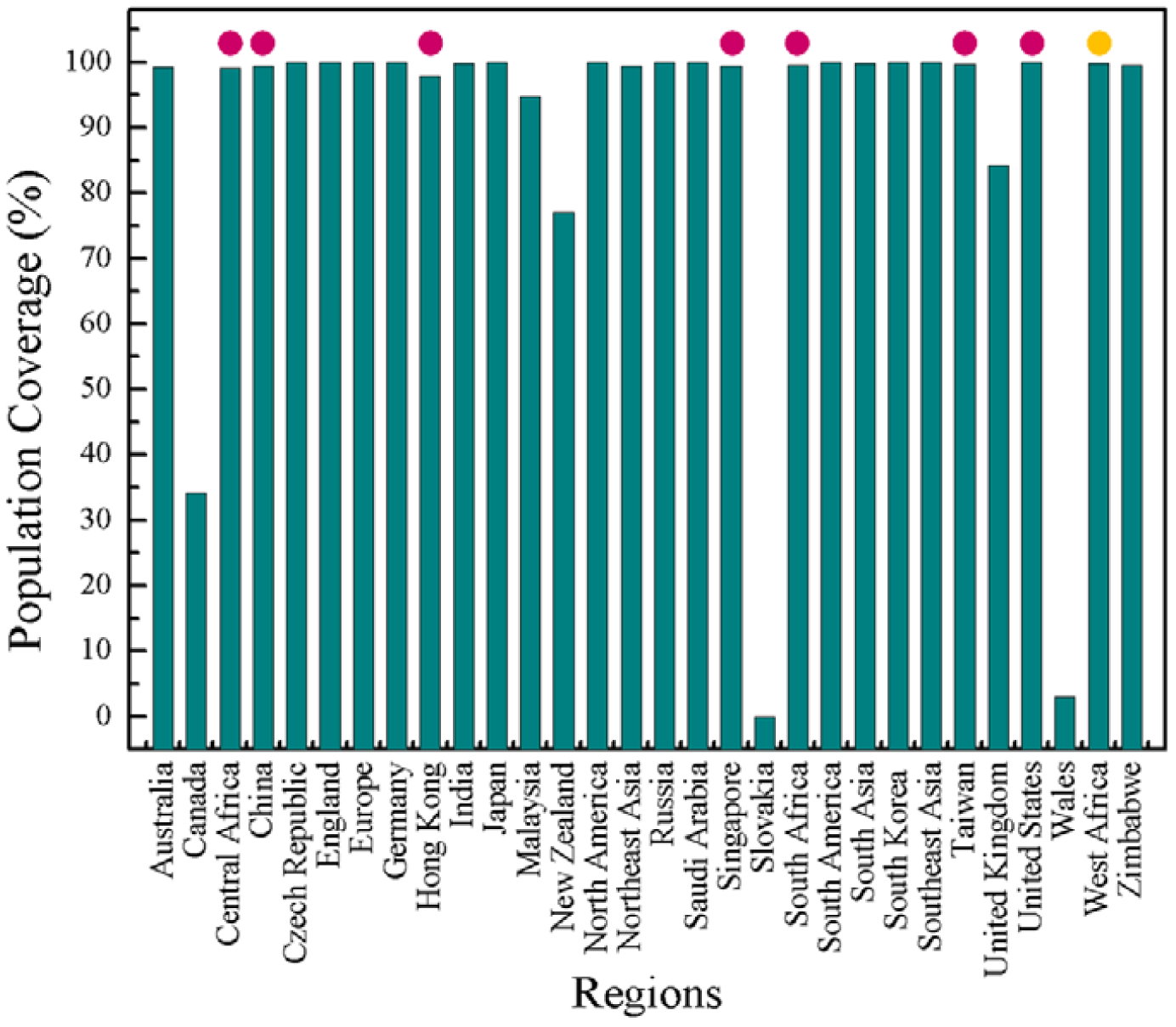
Worldwide population coverage by selected T-cell epitopes and their respective MHC binding alleles. Regions of particular importance were considered in this graph where pink and yellow circles are indicating the regions of outbreaks and bacterial origin, respectively.

### Molecular docking between epitopes and HLA alleles

In general, the epitopes having affinity to multiple alleles are better candidates for vaccine design because of high population coverage. Therefore, we considered 6 CTL and 7 HTL epitopes which have interactions to ≥5 of their respective alleles with an experimentally validated allele in common. We found that 6 CTL epitopes shared a common allele HLA-B*53:01, while 7 HTL epitopes shared an HLA-DRB5*01:01 allele as shown in Supplementary Table 5. So, these 13 epitopes (6 CTLs and 7 HTLs) were modelled using PEPFOLD 3 server. Finally, the molecular docking between these epitopic models and their respective common alleles was performed with AutoDock tools and AutoDock Vina. The binding affinities between epitopes and their binding alleles were compared to the control ligand which is given in Supplementary Table 6 and Figure 1. The best interacting CTL epitope model was LISLWFIIF (−10.0 kcal/mol) as compared to the CTL positive control (−9.3 kcal/mol) and the residues involved in classic hydrogen bond interactions were Leu1, Ile2, Ile7, Phe9, Asn63, Asn70, Asn77, Arg97, Tyr99, Thr143, Tyr123, Tyr9, Tyr171, Trp5, and Ser3 as shown in Fig. 3A. On the other hand, the HTL epitope model YFIWRAKENRFKHRQ was the best (−7.3 kcal/mol) as compared to HTL positive control (−8.5 kcal/mol). The conventional hydrogen bond network in YFIWRAKENRFKHRQ comprised of residue Gln9, Ser53, Glu55, Asn62, Tyr213, His281, Asn282, Arg5, Asn9, Arg14, Gln15, Glu8, His13, Phe2, Ala6, Phe51, and Asp270 which are shown in Fig. 3B. Thus, these 13 T-cell epitopes ensured their binding efficiency as well as their suitability to be used in multiepitope based vaccine design.

**Figure 3:**
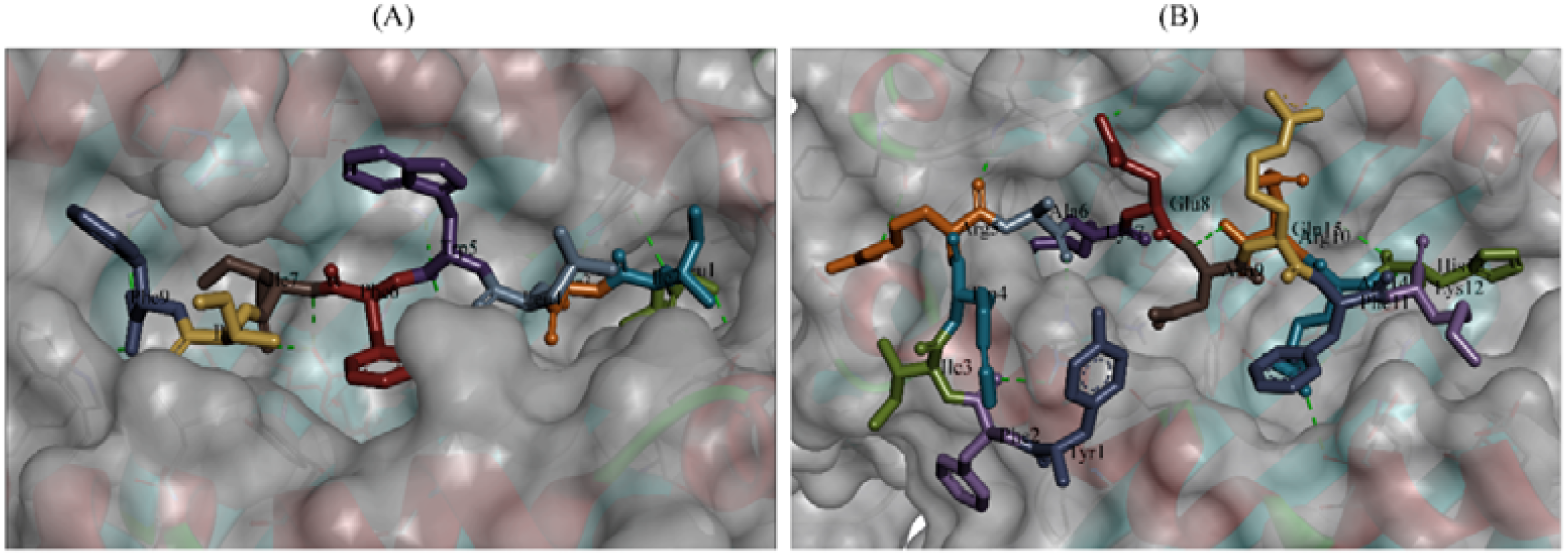
Molecular docking between T cell epitopes and their respective MHC alleles. (A) Molecular docking between best CTL epitope (LISLWFIIF) and HLA-B*53:01 allele where residue Leu1, Ile2, Ile7, Phe9, Asn63, Asn70, Asn77, Arg97, Tyr99, Thr143, Tyr123, Tyr9, Tyr171, Trp5, and Ser3 were involved in conventional hydrogen bonds, (B) Molecular docking between best HTL epitope (YFIWRAKENRFKHRQ) and HLA-DRB5*01:01 where residue Gln9, Ser53, Glu55, Asn62, Tyr213, His281, Asn282, Arg5, Asn9, Arg14, Gln15, Glu8, His13, Phe2, Ala6, Phe51, and Asp270 took part in conventional hydrogen bonds.

### Construction of multiepitope vaccine

In multiepitope vaccine design, we have considered highly potent T-cell epitopes that share at least one of their respective allele and B-cell epitopes that are antigenic, non-allergenic and non-toxic. Thus, a multiepitope vaccine is constructed using 7 HTL, 6 CTL and 2 BL epitopes fused by GPGPG, AAY, and KK linkers, respectively (Table 1 and 2). Epitopes in each group were sequenced according to their higher to lower antigenic score. Furthermore, TLR4 agonist 50S ribosomal protein L7/L12 (130 residues) was added as an adjuvant using EAAAK linkers. The arrangement of different epitopes along with their joining linkers is shown in Fig. 4. The final vaccine construct comprises 384 amino acid residues.

**Table 1:**
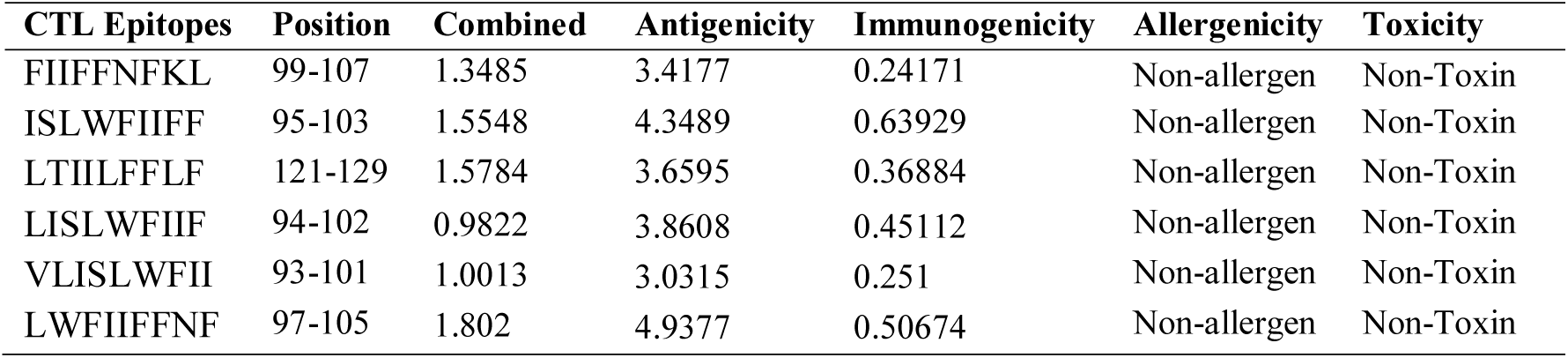
Final selected cytotoxic T lymphocyte (CTL) epitopes for multi-epitope vaccine construction.

**Table 2:**
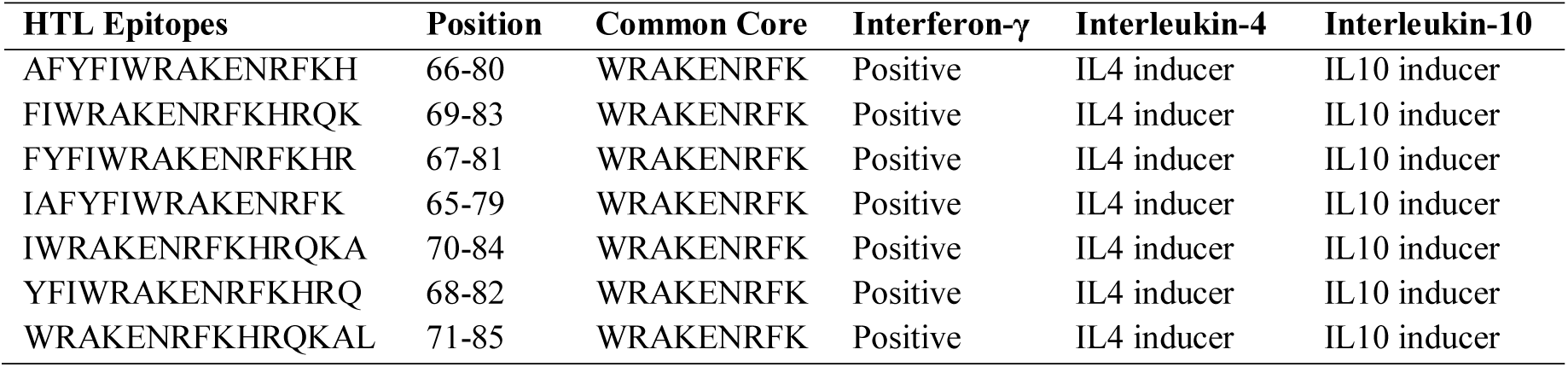
Final selected helper T lymphocyte (HTL) epitopes with their cytokine inducing properties.

**Figure 4:**
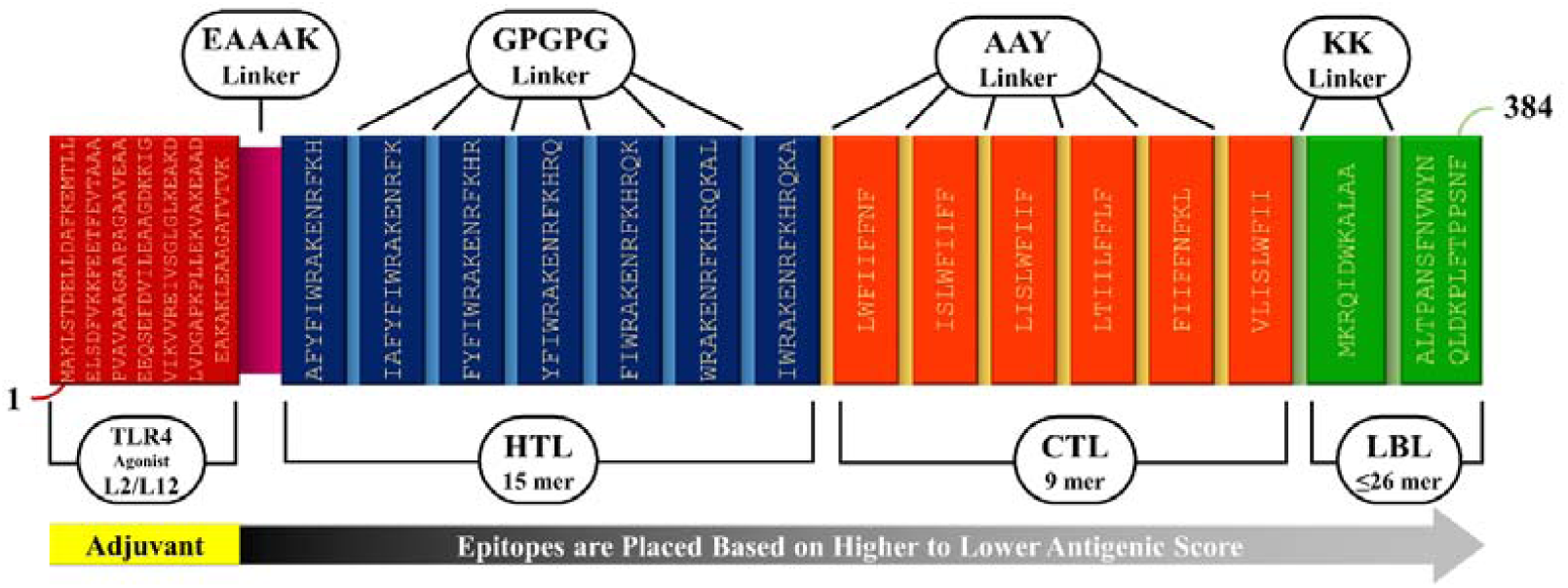
Graphical illustration of multi-epitope vaccine construct. The final vaccine construct is 384 amino acid residues long in which first 130 residues have been represented as adjuvant followed by HTL (131–269 amino acids), CTL (270–342 amino acids) and BCL (343-384) epitopes. Adjuvant and HTL epitope has been joined by EAAAK linker (maroon) while GPGPG (blue), AAY (yellow) and KK (light green) linkers were used to join the CTL, HTL and BL epitopes, respectively.

### Evaluation of the primary sequence of vaccine protein

The physiochemical and immunogenic nature of chimera protein was evaluated as shown in Table 3. The final vaccine construct was found to be basic, stable, thermostable, hydrophilic and highly soluble based on the analysis of physiochemical parameters. In addition, the half-life was estimated to be 30 hours in mammalian reticulocytes (*in vitro*), over 20 hours in yeast (*in vivo*) and over 10 hours in *E. coli* (*in vivo*). The immunogenic assessment also revealed that our vaccine construct is highly antigenic, immunogenic and non-allergenic (Table 3). These results indicated that this chimera protein is suitable to be a potential vaccine candidate.

**Table 3:** Antigenic, allergenic and physiochemical assessments of the primary sequence of final vaccine protein.

| Evaluating Parameters | Results | Remarks |
| --- | --- | --- |
| Number of amino acids | 384 | Suitable |
| Molecular weight | 43754.20 Dalton | Average |
| Chemical formula | C<sub>2079</sub>H<sub>3133</sub>N<sub>533</sub>O<sub>504</sub>S<sub>3</sub> | - |
| Theoretical pI | 9.99 | Basic |
| Extinction coefficient (at 280 nm in H<sub>2</sub>O) | 87890 M<sup>-1</sup>cm<sup>-1</sup> | - |
| Estimated half-life (mammalian reticulocytes, *in vitro*) | 30 hours | - |
| Estimated half-life (yeast cells, *in vivo*) | >20 hours | - |
| Estimated half-life (*Escherichia coli*, *in vivo*) | >30 hours | - |
| Instability index | 17.95 | Stable |
| Aliphatic index | 83.31 | Thermostable |
| Grand average of hydropathicity (GRAVY) | -0.076 | Hydrophilic |
| Antigenicity | 0.9258 | Antigenic |
| Immunogenicity | 7.034 | Immunogenic |
| Allergenicity | Probable non-allergen | Non-allergenic |
| Solubility | 0.72056 | Soluble |

### Secondary and tertiary structure prediction

The secondary structure was predicted using the latest PSIPRED 4.0 server which demonstrated the presence of 54.17% alpha helix (208), 8.59% beta strand (33) and 37.24% random coil (143) structure in our vaccine protein as depicted in Fig. 5A. Furthermore, homology modelling of vaccine protein was performed using the RaptorX server. A total of 384 amino acid residues was modelled as three domain(s) with 14% disorder (Fig. 5B). The RaptorX server uses multi-template based approach to build a protein model. For example, the template PDB ID: 1dd3A was chosen to model domain 1 (1-135 AA). Similarly, for domain 2 (232-384 AA) PDB ID: 5ujaA, 5u71A and 5tsiA; and domain 3 (136-231 AA) PDB ID: 5vflL were considered. P-value is a quality indicator in homology modelling and low P-value defines the high-quality model. The *P*-value obtained for the modelled structure was 8.67e-04, which is low and significant (Supplementary Table 7).

**Figure 5:**
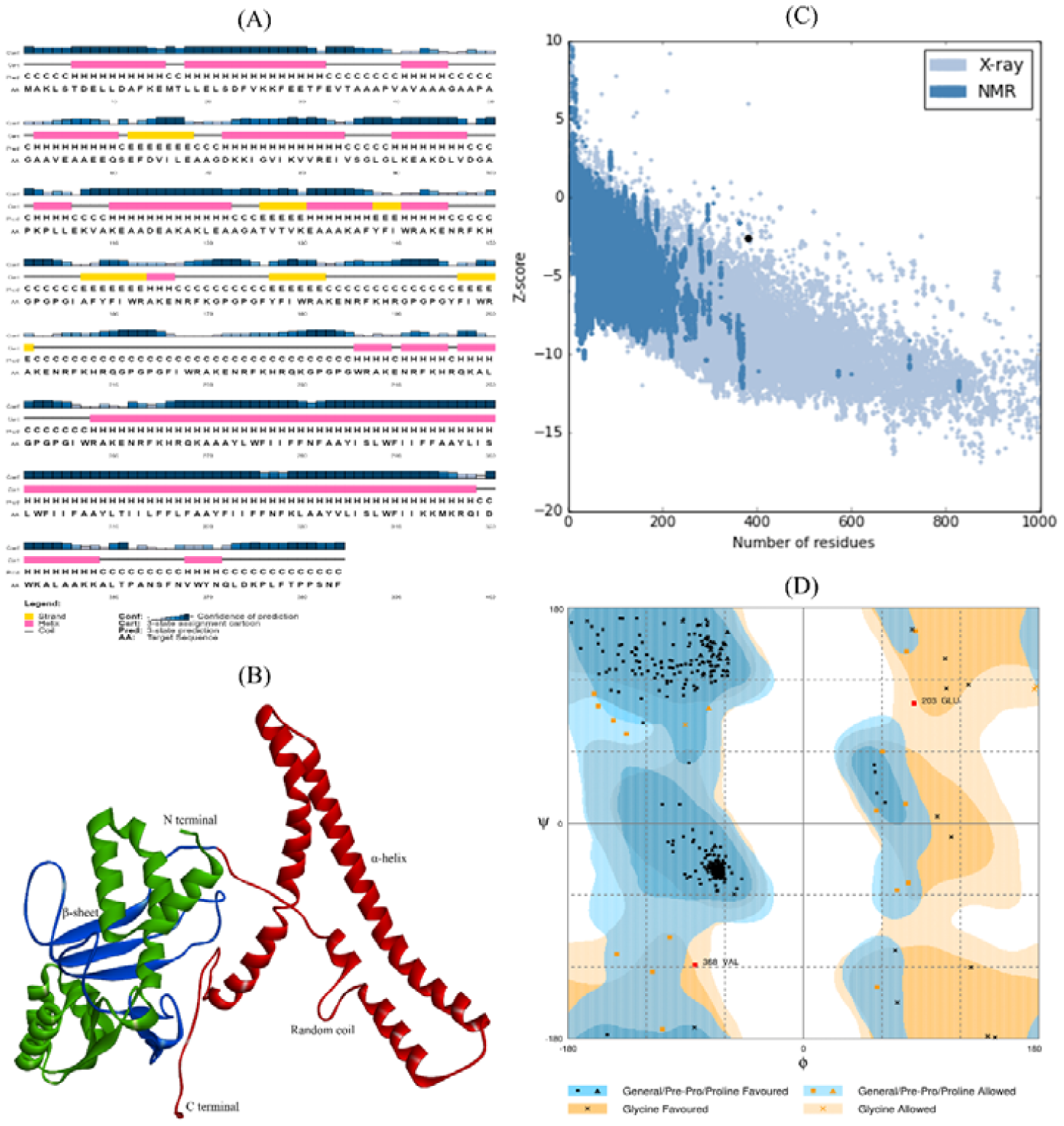
Prediction and assessment of the secondary and tertiary structure of the vaccine construct. (A) cartooned secondary structure of final vaccine construct showing α-helix (54.17%), β-strands (8.59%) and coils (37.24%), (B) 3D structure of vaccine protein showing three domains in different colours, (C) ProSA validation of 3D vaccine model showing Z-score (−2.54), and (d) Ramachandran plot analysis of refined structure showing 94.2%, 5.2% and 0.5% residues in favoured, allowed and disallowed region, respectively.

### Refinement and validation of tertiary structure

Structure refinement with GalaxyRefine leads to an increase in the number of residues in the favoured region. Of all the refined models, the model 3 proved to be the best on the basis of various parameters including GDT-HA (0.9206), RMSD (0.527), MolProbity (2.502), Clash score (23.9), Poor rotamers (2.0) and Rama favoured (94.0) as shown in Supplementary Table 8. This model was taken as the final vaccine model and subjected to structural validation by plotting a Ramachandran plot and found the same that 94.2% (360) residue in Rama-favoured region, 5.2% (20) residues in allowed region and only 0.5% (2) residues in outlier region (Fig. 5D). ProSA-web verified the quality and potential errors in a crude 3D model. The overall quality factor of a modelled protein was 82.2% using ERRAT. While ProSA-web has shown the Z-score of −2.54 for the input vaccine protein model which is lying outside the scores range that commonly found in the case of native proteins of comparable size (Fig. 5C).

### Mapping of B-cell epitopes in vaccine construct

The presence of B-cell epitopes was predicted using the ElliPro tool of IEBD, allowing the identification of 9 linear B-cell epitopes with a score of 0.527 to 0.819 and 10 conformational B-cell epitopes comprising of a total of 205 amino acid residues as shown in Fig. 6A and Fig. 6B, respectively. The scores of the conformational B-cell epitopes ranged from 0.536 to 0.920, while the size ranged from 3 to 64 residues.

**Figure 6:**
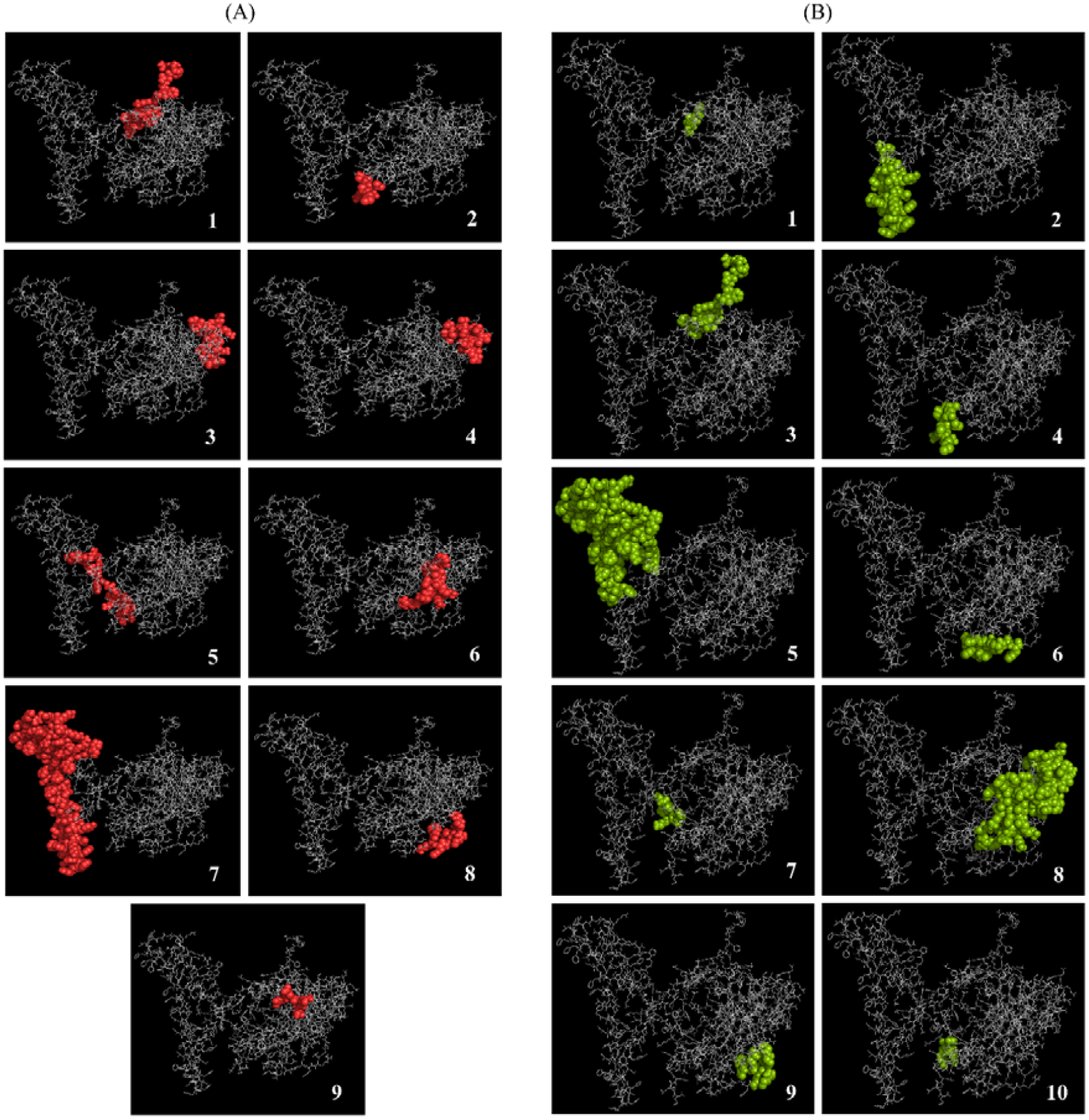
B lymphocyte epitope present in the vaccine: (A) Red spheres showing linear BL epitopes containing 1. 24 residues with 0.819, 2. 5 residues with 0.803, 3. 16 residues with 0.655, 4. 14 residues with 0.654, 5. 19 residues with 0.552, 6. 11 residues with 0.543, 7. 85 residues with 0.805, 8. 8 residues with 0.628 and 9. 4 residues with 0.527 score); and (B) Green spheres showing conformational BL epitopes containing 1. 3 residues with 0.92, 2. 30 residues with 0.833, 3. 21 residues with 0.805, 4. 6 residues with 0.78, 5. 64 residues with 0.743, 6. 10 residues with 0.671, 7. 7 residues with 0.633, 8. 59 residues with 0.59, 9. 7 residues with 0.56, and 10. 3 residues with 0.532 score).

### Disulphide bridging for vaccine stability

A total of 35 pairs of residues were found that could be used for disulphide engineering which is given in Supplementary Table 9. However, after evaluating other parameters such as energy and χ3, only a pair of residuals were finalized because their value falls below the allowable range *i*.*e*., the value of energy should be less than 2.2 kcal/mol and χ3 angle should be in between −87 and +97 degree^37^. Therefore, a total of two mutations were generated on the residue pairs, Gly173-Glu203, for which the χ3 angle and the energy were −86.78 degree and 1.91 kcal/mol, respectively as pointed in Fig. 7.

**Figure 7:**
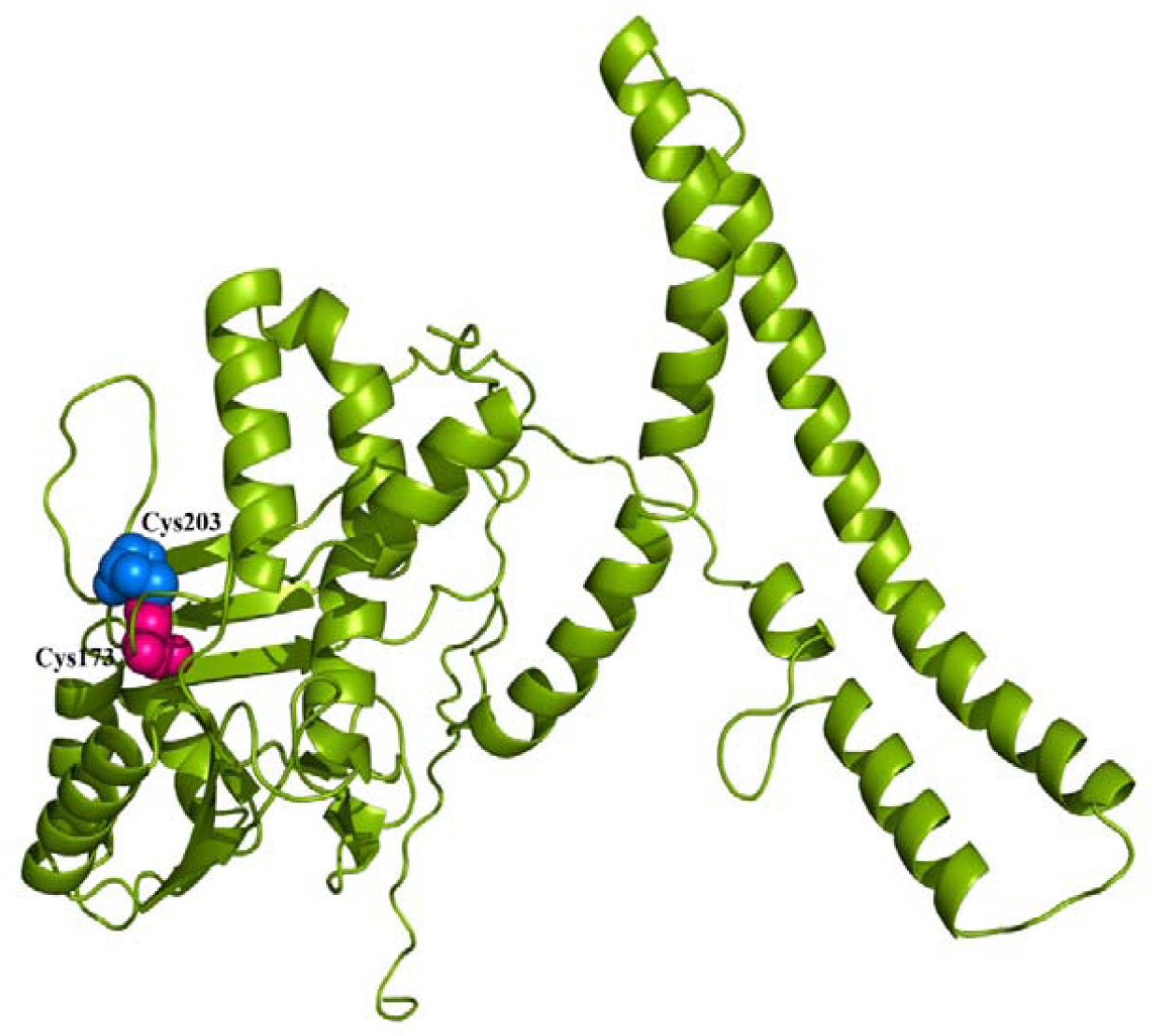
Disulphide engineering of the final vaccine protein. Mutated residue pairs showed in pink (initial residue: Gly173) and blue colour (initial residue: Glu203). Residues were selected based on their energy, χ3 value, and B-factor.

### Codon adaptation and *in silico* cloning

The main purpose of *in silico* cloning was to express the *E. anophelis*-derived vaccine protein epitope into the *E. coli* expression system. Therefore, it was necessary to adapt the codon respective to the subunit vaccine construct as per the codon usage of *E. coli* expression system. We adapted the codons according to the *E. coli* K12 strain using JCAT server and found that the GC-content of the improved sequence was 49.39% while the value of the codon adaptive index was 0.95 close to 1.0. Therefore, a satisfactory adaptation was achieved. Later on, XhoI and NdeI restriction sites were created and cloned into the vector pET28a (+) as shown in Fig. 8. The target sequence is also included at both ends between 6-histidine tag residues, which is helpful for purification purposes. Thus, the total length of the clone was 6.45 kbp.

**Figure 8:**
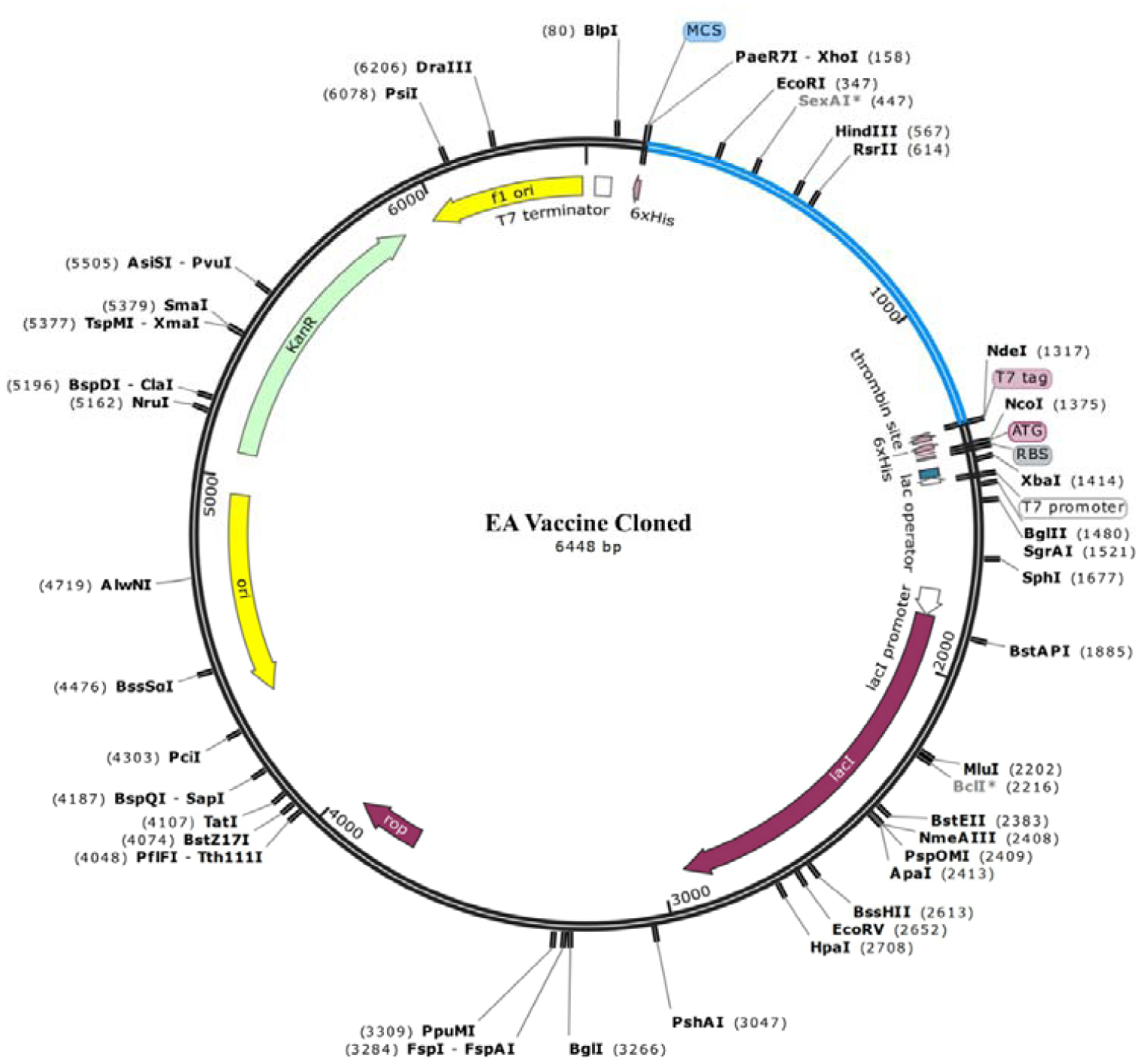
*In silico* cloning. Vaccine protein sequence adapted to pET28a (+) vector showing the region of choice in blue colour surrounded between XhoI (158) and NdeI (1317) while the vector has shown in black lines.

### Molecular docking of vaccine with TLR4 receptor

A total of 30 models of the vaccine-receptor complex was generated in molecular docking. Energy scores for these models are tabulated in Supplementary Table 10. Among these occupied models, the receptor was appropriately located and had the lowest energy value for model number 5, which was therefore selected as the best-docked complex (Fig. 9A). The energy score obtained for model 5 was found to be −1626.2 which is lowest among all predicted docked complexes and thus, shows the highest binding interaction.

**Figure 9:**
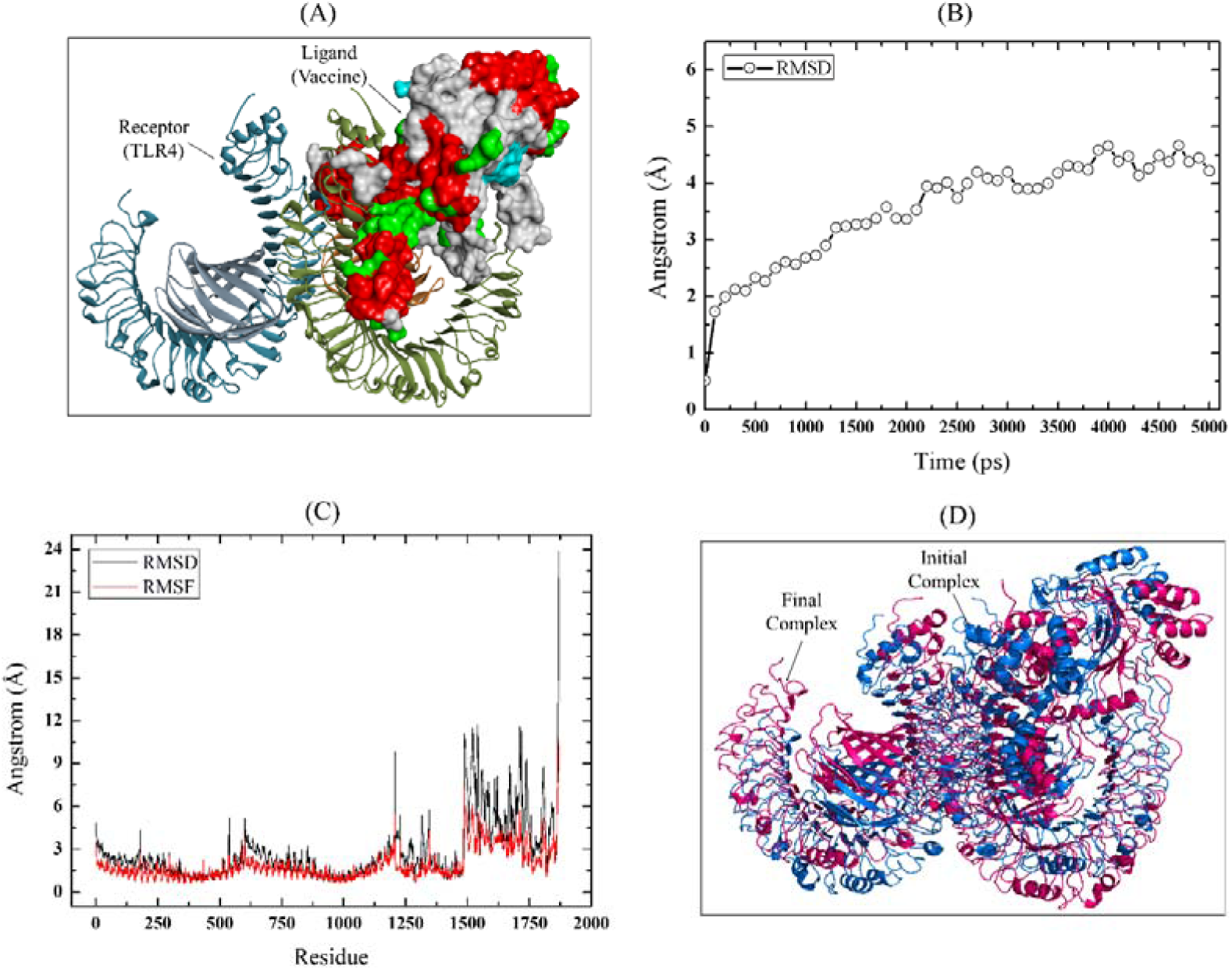
Molecular docking and dynamic simulation studies. (A) Molecular docking between vaccine (sphere) and TLR4 receptor, (B) RMSD value of the vaccine-TLR4 complex backbone; (C) Plot showing the RMSF (red) and RMSD (black) values of side-chain residues, and (D) Conformational illustration of initial (blue) and final (pink) step of vaccine-TLR4 complex in 5 ns long dynamic simulation.

### Dynamic simulation of the vaccine-TLR4 complex

The vaccine-TLR4 complex becomes stabilized around 4 ns in the 5 ns long dynamic simulation as shown in Fig. 9B. The average simulation energy was −7261872.45 kJ/mol, while the Coulombic charge and van der Waals interactions were −9968104.77 and 1284708.44 kJ/mol, respectively (Supplementary Table 11). The RMSD was calculated to determine the structural stability of the vaccine-TLR4 complex. The average RMSD of the complex backbone was 3.55 Å while the RMSD value of side-chain residue varied between 0.73 Å to 24.44 Å (Fig. 9C). Furthermore, RMSF was observed to evaluate the side-chain flexibility. The RMSF value ranged from 0.61 Å to 10.68 Å and the higher peaks at ∼10.68 Å in the plot indicate highly flexible regions in the complex (Fig. 9C). The illustration of the initial and final state of the complex demonstrated that both vaccine and TLR4 receptor have remarkable changes in their conformation to facilitate the binding to form a stable complex in the thermodynamic environment (Fig. 9D). In addition, the number of initial and final hydrogen bonds were 1019 and 675, respectively. On average, 650 hydrogen bonds were present during the simulation, suggesting strong binding of the vaccine-receptor complex. These values indicate strong complex stability based on previous observations^34,35^. Slight fluctuations in amino acid side chains also indicate the uninterrupted interaction between vaccine and receptor.

### *In silico* immune response simulation

The *in silico* immune response generated in C-ImmSim immune simulator was consistent with actual immune responses as depicted in Fig. 10. The secondary and tertiary responses were significantly higher than the primary response. The normal high levels of immunoglobulin activity (*i*.*e*., IgG1 + IgG2, IgM, and IgG + IgM antibodies) was evident in both secondary and tertiary responses with a corresponding decrease in antigen concentration. In addition, several B-cell isotypes with long-lasting activity were observed, indicating possible isotype switching potentials and memory formation (Fig. 10A-C). A similarly high response was seen in the TH (helper) and TC (cytotoxic) cell populations with corresponding memory development (Fig. 10D-F) which is essential for complementing the immune response. During exposure, increased macrophage activity was demonstrated while dendritic cell activity was found to be consistent (Fig. 10G-H). High levels of IFN-γ and IL-2 were also evident. In addition, a lower Simpson index (D) indicates greater diversity (Fig. 10I). Moreover, repeated exposure to 12 injections elicited increasing IgG1 and decreasing IgM levels, while maintaining IFN-γ and TH cell populations at a high level throughout exposure as provided in Supplementary Figure 2. This profile indicates the development of immune memory and, consequently the increased clearance of the antigen at subsequent exposures.

**Figure 10:**
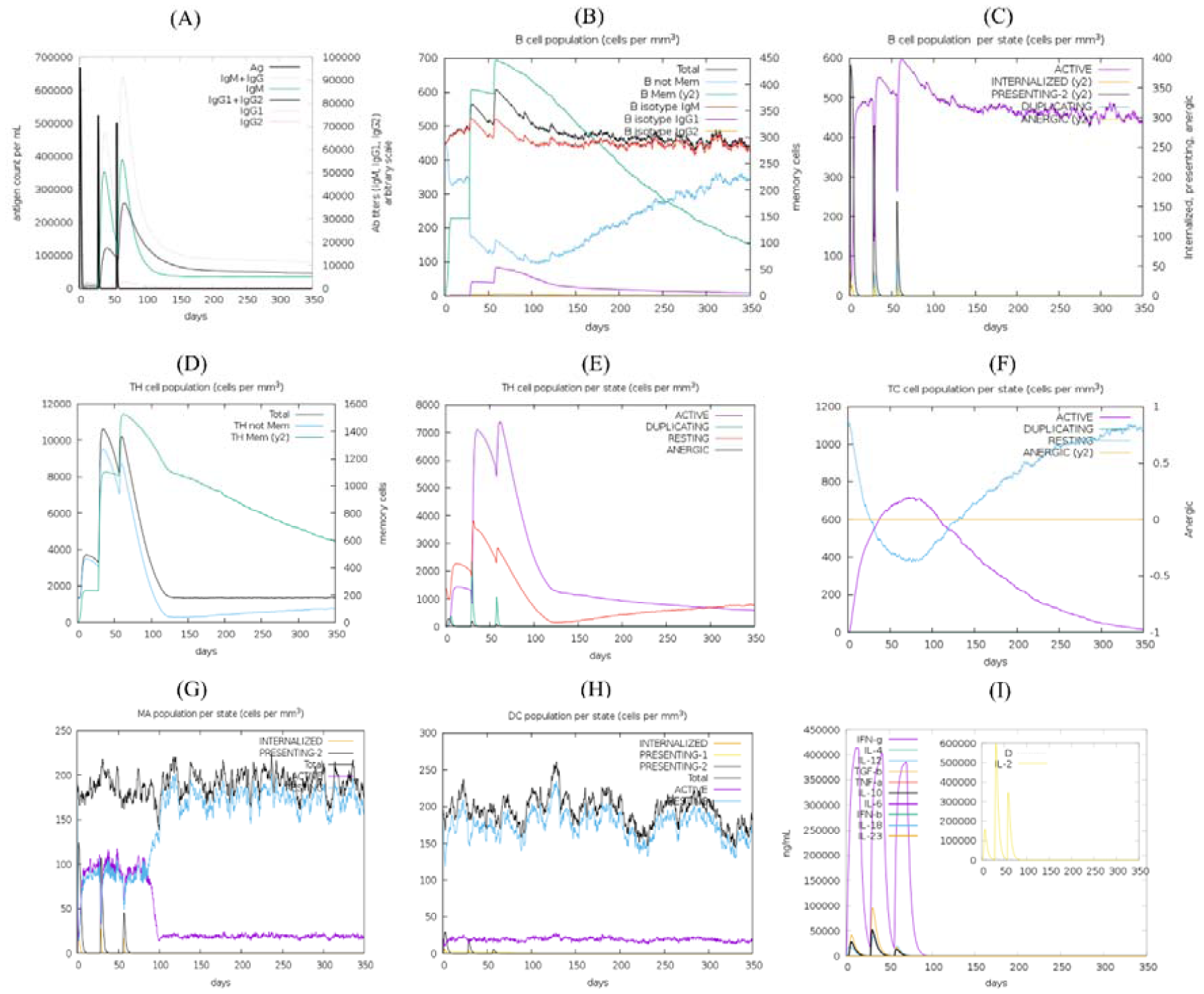
*In silico* simulation of immune response using vaccine as antigen: (A) Antigen and immunoglobulins, (B) B cell population, (C) B cell population per state, (D) TH cell population, (E) TH cell population per state, (F) TC cell population per state, (G) MA population per state, (H) DC population per state, and (I) Production of cytokine and interleukins with Simpson index (D).

## Discussion

Currently, vaccines are the most promising mode that improves the immune system^27^. However, the development and production of the vaccine are labour-intensive and costly^38^. Immunoinformatics can reduce this burden where a variety of tools and servers are applied to identify potential B and T-cell epitopes, which subsequently resulted in the designing of epitope-based subunit vaccine. Despite the recent outbreaks worldwide, there is no preventive measure that can suppress *E. anophelis*. In this study, the translocator protein (TSPO) in the *E. anophelis* proteome showed the highest antigenicity. TSPO belongs to a membrane protein family (Pfam: TspO/MBR family PF03073) and is associated with stress, membrane transport, and some neurodegenerative diseases^39^. Therefore, the higher antigenic nature of this protein is no wonder in this case. Furthermore, we found that *E. anophelis*-derived TSPO protein is not homologous to its human counterpart or any other proteins of human origin. Therefore, TSPO is a perfect target for the design and testing of a multiepitope vaccine against *E. anophelis*.

The final vaccine protein comprised of antigenic, non-allergenic, and non-toxic B-cell and T-cell epitopes with higher binding affinity for several MHC molecules, which in turn provided high population coverage (99.99%) worldwide. It is important that it covers more than 97.5% population of *E. anophelis* outbreak regions. Besides, the highly antigenic and non-allergenic property of the final vaccine protein make it a potent vaccine construct. The molecular weight of vaccine protein was 54.62 kDa which is an average molecular weight for a multisubunit vaccine. Further, the theoretical pI indicates that our vaccine protein is basic and is most stable in the pH range of 9.04. Furthermore, the aliphatic index and instability index classify our vaccine as thermostable, while the negative GRAVY value suggests that the vaccine protein is hydrophilic and has strong interactions with water molecules. However, one of the biggest drawbacks in the field of therapeutic proteins is the short half-life of peptides^40^. Interestingly, our vaccine protein half-life in mammalian reticulocytes (*in vitro)* was 30 hours and more than 20 hours and 10 hours in yeast and *E. coli* cells *in vivo*, respectively which is satisfactory based on earlier findings^34,35^. Also, the recombinant proteins should be soluble on overexpression for post-production studies^41^. The predicted solubility of the vaccine protein ensured their ease access.

The tertiary structure of the vaccine was predicted and refined, followed by validation. The refinement of the crude model was ensured by their parametric scores. For instance, the crude model had 92.4% residues in Rama favoured zone while refined models have up to 94.8% (Supplementary Table 8). In addition, GDT-HA score between 0.9 and 1.0 suggested great similarity of the refined models with the amino acid sequence. The Z-score (−2.54) and ERRAT quality factor (82.196) indicate that the overall quality of the refined vaccine structure was satisfactory. Besides, the Ramachandran plot shows that 94.2% of residues clustered tightly in the most favoured region with very few residues in outliers. A good quality model would probably exceed 90% in the most favoured regions^42^. Hence, the 3D structure of the vaccine is considered good. The expression of our recombinant vaccine protein inside *E. coli* K12 was improved through codon adaptation. The codon adaptation index (CAI) and GC content of the optimized DNA sequence were 0.95 and 49%, respectively. The CAI value (0.95) closer to 1.0 suggests the higher expression probability. The vaccine protein should be mechanically stable which is accomplished by creating unique disulphide bonds into multisubunit vaccine construct.

Further, molecular docking and dynamic simulation were performed to ensure the binding efficiency and stability of the vaccine-receptor complex, respectively. It is clear that the vaccine protein could properly occupy the TLR4 receptor with minimal energy. The simulated microscale-change in protein backbone (RMSD: 3.55 Å) and mild fluctuations in side-chain residues confirmed the stability of the vaccine-TLR4 complex. In addition, binding interactions within the complex were good according to hydrogen bond network analysis. Finally, *in silico* immune response simulation showed higher B and T-cell activity (helper T-cells in particular), and the results were in coherence with typical immune responses. The formation of B and T-cell memory was evident and last for several months. The high level of IFN-γ and IL-2 production during repeated exposure confirms the efficient Ig production, thereby, supporting a humoral response. Importantly, macrophages and dendritic cells are important when the expression of TLR4 in human is considered^43^. In our study, both macrophage and dendritic cell activities were satisfactory. Moreover, components of the innate immune system such as epithelial cells were also active. The increase in Simpson index (D) over time indicates the emergence of a dominating clone that means the higher the value, the lower the diversity^44^. Therefore, the lower D value suggests a diverse immune response that is plausible considering the generated chimeric peptide which is composed of several B and T-cell epitopes.

However, there are certain limitations in immunoinformatics approach as well. For example, lack of standard benchmark, inadequate prediction methods, and unavailability of precise datasets for different computational analysis^32^. In addition, we have no idea about the exact receptor and path used by *E. anophelis* for their pathogenic entry, along with their specific immune-bypass mechanism within the human host. With this knowledge, the vaccine could be developed more effectively and efficiently to control and/or prevent the pathogenesis of *E. anophelis*.

## Conclusion

*E. anophelis* has emerged as an opportunistic human pathogen involved in neonatal meningitis, sepsis, and several major nosocomial outbreaks. Despite their significant causality rate and antimicrobial resistance profile, the transmission route and control measure of this pathogen remain unattainable. In this study, we have utilized immunoinformatic approaches along with molecular dynamics and immune simulations to generate an effective multiepitope vaccine candidate. Therefore, the highest antigenic protein from *E. anophelis* proteome was selected followed by the prediction of T-cell and B-cell epitopes to produce cell-mediated and humoral immunity, respectively. Predicted epitopes were merged using suitable linkers and adjuvant to enhance the immunogenicity and adequate separation of epitopes within the human body. Allergenicity and antigenicity were also confirmed followed by the physiochemical properties evaluation. Molecular docking and dynamics were also performed to check the binding affinity and stability of TLR-4 and vaccine complex. Finally, disulphide engineering and *in silico* cloning were introduced to ensure the stability and effective expression of the vaccine construct. However, the proposed vaccine needs to be experimentally validated to ensure the control of *E. anophelis* infection by producing an active immunological memory.

## Methodology

### Retrieval of protein sequences and prediction of the highest antigenic protein

The FASTA formatted whole proteome of the *Elizabethkingia anophelis* Ag1, except protein homologs in *Homo sapiens*, was retrieved from the Integrated Microbial Genomes & Microbiomes 5.0 (IMG/M) system 45. It is heavily resourced by NCBI database^46^ and integrates publicly available draft and complete genomes from all three domains of life^47^. The ‘Phylogenetic Profiler’ tool of IMG/M system allows the identification of protein sequence for each gene in a query genome with or without the protein homologs in other genomes^47^. Antigenicity is the ability to induce an immune response towards an antigen. Therefore, the selection of proteins with higher antigenicity is a prerequisite for peptide-based vaccine design. So, all proteins from *E. anophelis* proteome were evaluated using the VaxiJen 2.0 server with 0.5 threshold. This server maintains 70-89% prediction accuracy through auto cross-covariance (ACC) transformation^48^.

### Prediction of cytotoxic T lymphocyte (CTL) epitopes and their MHC-I binding alleles

Most cytotoxic T-cells express T-cell receptors (TCRs) that can recognize a specific antigen^49^. Therefore, prediction of CTL epitopes is essential for coherent vaccine design. The selected protein was submitted to the NetCTL 1.2 server for the prediction of CTL epitopes with 0.5 threshold^50^. This server predicts CTL epitopes based on three criteria, namely, MHC-I binding peptides, C terminal cleavage, and TAP transport efficiency. MHC-I binding and C-terminal cleavage are obtained by using artificial neural networks whereas TAP transporter efficiency is evaluated by the weight matrix 50. Furthermore, MHC-I binding alleles for each 9-mer CTL epitope were predicted using the consensus method in the IEDB analysis tool^51^. The MHC source species was human and percentile rank ≤2 was considered.

### Assessment of CTL epitopes for immunogenicity, allergenicity, and toxicity

The suitability of CTL epitopes lies within their immunogenic potential, hence, considered as the centre of the vaccine adeptness^26^. So, all predicted CTL epitopes were evaluated for their immunogenicity using MHC-I immunogenicity tool of IEDB server^52^. Further, their antigenicity is evaluated again to ensure their ability to induce immune response through VaxiJen 2.0 server^48^. Vaccine components should be free from allergic reaction. So, AllerTOP 2.0 server was used for allergenicity prediction. This server is designed to predict allergenicity based on κ-nearest neighbours (kNN) method with 88.7% accuracy^53^. Furthermore, epitopes having toxicity should be eliminated since they could compromise the aim and development of the vaccine. Therefore, we used ToxinPred server to identify the toxic CTL epitopes. This server works based on the different properties of peptides using machine learning technique and quantitative matrix^54^.

### Prediction of helper T lymphocyte (HTL) epitopes and their MHC-II binding alleles

Helper T-lymphocytes (HTLs) are a part of the adaptive immune response and can generate cellular and humoral immune response against pathogens or foreign antigens. Therefore, the prediction of HTL epitopes which bind to MHC-II alleles are essential and play a significant role in rational vaccine design^55^. The HTL epitopes of 15-mer length (peptide) and their corresponding MHC class II alleles were predicted through the IEDB MHC II binding tool using the consensus method^56^. For the consensus method, a percentile rank is generated by comparing the peptide’s IC50 against those of a set of random peptides from the SWISS-PROT database^57^. A lower percentile rank indicates high affinity, therefore, percentile rank ≤2 was considered in this study.

### Identification of cytokine-inducing HTL epitopes

The helper T-cells help in the activation of cytotoxic T-cells and other immune cells upon releasing different types of cytokines *i*.*e*., interferon-gamma (IFN-γ), interleukin-4 (IL-4) and interleukin-10 (IL-10)^58^. As a result, cytokine-inducing HTL epitopes are crucial for the development of vaccine or immunotherapy. Therefore, we used IFNepitope server to predict IFN-γ producing HTL epitopes with Motif and SVM based hybrid method and IFN-γ versus Non-IFN-γ model as the default parameters^59^. On the other hand, IL-4 and IL-10 inducing properties were predicted using IL4pred^60^ and IL10pred^61^ servers, respectively. These two servers are designed for *in silico* prediction of interleukin inducing peptides. The IL4pred and IL10pred operations were carried out based on SVM method and hybrid method (Motif and SVM), respectively with other parameters kept as default.

### Prediction and assessment of linear B lymphocyte (LBL) epitopes

B-cells provide humoral immunity by secreting immunoglobulins which can neutralize antigen upon binding^62^. Hence, B-cell epitopes are also essential for effective vaccine construct. B-cell epitopes are two types, linear (continuous) and conformational (discontinuous). The ElliPro tool from the IEDB database can predict both types of B-cell epitopes based on flexibility and solvent-accessibility^63^. In case of vaccine design, however, only the linear epitopes are considered. Therefore, only LBL epitopes were predicted using the ElliPro tool with default parameters. The tool utilizes three different algorithms (*i*.*e*., an approximation of protein shape, protrusion index (PI) of residues and clustering of neighbouring residues) where each predicted epitope was given a PI score. Residues with larger PI values are related to greater solvent accessibility^63^. The predicted LBL epitopes were subjected to further evaluation in terms of their antigenic, allergenic and toxic nature using VaxiJen 2.0, AllerTOP 2.0 and ToxinPred server, respectively.

### Estimation of population coverage

The distribution of HLA alleles, as well as their expression, could vary throughout the world according to the difference in regions and ethnicities^25^. Therefore, successful vaccine development demands the evaluation of HLA allele distribution across the world population^64^. In this study, the distribution of HLA alleles for potential CTL and HTL epitopes was evaluated with the IEDB population coverage tool^64^. This server is designed to estimate the population coverage of epitopes from different regions based on the distribution of their MHC binding alleles. Since the prevalence of *E. anophelis* is poorly understood, the percentage of population coverage was calculated for different parts of the world. In particular, however, we emphasized the areas where it was first appeared and caused outbreaks.

### Peptide modelling and molecular docking studies

Epitopes having a common corresponding allele with good binding interactions to more alleles are good choice for multiepitope vaccine design. Therefore, epitopes showing binding interaction to an experimentally validated common allele were targeted for molecular docking studies using the AutoDock tools^65^ and AutoDock Vina software^66^. Molecular docking is a way to measure the binding affinity between protein and ligand molecule. At first, the selected CTL and HTL epitopes were modeled using PEP-FOLD 3.0 server with sOPEP sorting scheme in 200 simulations^67^. This server was designed to predict the conformations of small peptides (5-50 amino acids) based on Forward Backtrack/Taboo Sampling algorithm^67^. For CTL epitopes, the most conserved allele HLA-B*53:01 protein (PDB ID: 1A1O) was downloaded from RCSB Protein Data Bank (PDB). The allele was then processed with Discovery Studio and AutoDock tools. The grid box centre was set at 5.78, 28.96 and 21.71 Å in the X, Y and Z axis, respectively while the size was set at 54, 42 and 28 Å in the X, Y and Z dimensions, respectively. Similarly, the allele HLA-DRB5*01:01 (PDB ID: 1HQR) was considered for HTL epitopes where grid box centre was set at 18.72, 27.28 and 28.80 Å in the X, Y and Z axis, respectively, and the size was set at 44, 38, and 34 Å in the X, Y and Z dimensions, respectively. The spacing was 1.0 Å for both cases. Finally, the docking simulation was performed using AutoDock Vina. The PyMOL molecular graphics system (PyMOL) was used to make docked complex and visualized in BIOVIA Discovery Studio 2017.

### Designing of multiepitope vaccine construct

The multiepitope vaccine construct was designed by combining adjuvant, CTL, HTL and LBL epitopes with appropriate linkers as reported earlier^27,36^. An adjuvant is an immunogenic composition that may boost the specific and long-lasting immune response against an antigen by a variety of mechanisms^68^. Toll-like receptor (TLR) agonists are well-known for recognizing ligands from many Gram-negative bacteria and also to be a part of the peptide-based subunit vaccine^35^. In addition, TLR4 receptor upregulates TSPO/MBR related proteins by overproducing cytokines^69^. Therefore, a TLR4 agonist namely 50S ribosomal protein L7/L12 was retrieved from the NCBI database (Accession no. P9WHE3) and used as an adjuvant. For the effective separation of each epitope within the human body, EAAAK, GPGPG, AAY, and KK linkers were used. EAAAK linker was designed for effective separation of bifunctional fusion protein domains^70^, therefore, utilized to join the adjuvant and first HTL epitope. For efficient recognition of multiepitope vaccine, GPGPG and AAY linkers were used to merge HTL and CTL epitopes. The bi-lysine (KK) linker was used to fuse B-cell epitopes to preserve their independent immunogenic activities^71^.

### Immunogenic, allergenic and physiochemical evaluation of vaccine construct

Immunogenicity is the ability to stimulate cellular and humoral immune response whereas antigenicity is the ability to recognize specific antigen followed by the immune response. In other words, all immunogens are antigens, but not all antigens are immunogens^72^. Therefore, vaccine candidate should be highly antigenic as well as immunogenic in nature. Consequently, the predicted vaccine construct was evaluated with VaxiJen 2.0 server^48^ and IEDB Immunogenicity tool^52^ to ensure its antigenic and immunogenic potentials, respectively. On the contrary, allergenicity is the ability to create sensitization and allergic reactions associated with the IgE antibody response. So, vaccine candidate must be non-allergenic. We used AllerTOP 2.0 server to check the allergenicity of vaccine construct. Various physiochemical features of our vaccine protein *i*.*e*., theoretical pI, instability index, in vitro and in vivo half-life, aliphatic index, molecular weight and grand average of hydropathicity (GRAVY) were assessed by using ProtParam server^73^. Furthermore, the solubility of the vaccine protein upon overexpression in *E. coli* was predicted by SOLpro tool which has over 74% accuracy with 10-fold cross-validation^41^.

### Prediction of secondary and tertiary structures

The primary sequence of vaccine protein was submitted to PSIPRED 4.0 server to predict the secondary structure. PSIPRED server uses two feed-forward neural networks which process the PSI-BLAST (Position-Specific Iterated–BLAST) generated a position-specific scoring matrix to predict a precise secondary structure from the input sequence^74^. The normal Q_3_ score of an earlier version (version 3.2) of PSIPRED server was 81.6%^34^. The tertiary structure of a protein provides maximum stability in its lowest energy state through proper bending and twisting. Therefore, we used the RaptorX server to predict the 3D structure of our final vaccine construct^75^. RaptorX server predicts 3D protein model based on multiple-template threading (MTT) and provides some confidence scores to indicate the quality of a predicted 3D model. For example, P-value for relative global quality, GDT, and uGDT for absolute global quality, and modelling error at each residue^75^. The obtained 3D structure was visualized with BIOVIA Discovery Studio 2017.

### Refinement and validation of 3D vaccine model

The crude model of our multiepitope vaccine protein was refined using GalaxyRefine web-server^76^ which is based on the CASP10 tested refinement method^77^. GalaxyRefine performs rehashed structure perturbation followed by overall structural relaxation by performing molecular dynamics simulation^35^. Structure validation is an important step in homology modelling which works based on experimentally validated 3D protein structure. Hence, the refined vaccine model was submitted to ProSA-web for structural validation^78^. ProSA calculates an overall quality score for a specific input structure. The quality score outside the characteristic range of native proteins indicates possible errors in the predicted protein structure. ERRAT server^79^ was used to analyse the statistics of non-bonded interactions. Ramachandran plot was created using the RAMPAGE server^80^. This server validates protein structure in terms of energetically allowed and disallowed dihedral angles psi (ψ) and phi (□) of amino acid on the basis of PROCHECK principle^81^.

### Prediction of B-cell epitopes in the vaccine protein

B lymphocyte provides humoral immunity by producing antibodies. Besides, they present antigen and secrete cytokines^82^. Therefore, the vaccine construct should have B-cell epitopes within their protein domains. As we mentioned earlier, B-cell epitopes can be divided into two categories: linear (continuous) and conformational (discontinuous). The presence of both linear and conformational B-cell epitopes was determined by using the ElliPro tool of IEDB server^63^.

### Disulphide engineering of the final vaccine

Disulphide engineering is a novel approach for creating disulphide bonds into the target protein structure. Disulphide bonds are covalent interactions that help in increasing the protein stability along with the examination of protein interactions and dynamics^34,35^. Therefore, the refined model of the final vaccine construct was subjected to the Disulphide by Design 2.12^37^ web-platform to perform disulphide engineering. Initially, the refined protein model was uploaded and run for the residue pair search that can be used for the disulphide engineering purpose. Potential residue pairs were selected for mutation with cysteine residue using create mutate function of the Disulfide by Design 2.12 server.

### *In silico* codon adaptation and cloning

Codon adaptation is a way to increase the translational efficiency of foreign genes in the host when codon usage in both organisms differs from each other. In other words, unadapted codon may lead to the minor expression rate in the host. Therefore, the vaccine protein sequence was submitted to the Java Codon Adaptation Tool (JCAT)^83^ to adapt their codon usage to the most sequenced prokaryotic organism, *E. coli* K12^34^. Three additional options were selected to avoid the rho-independent transcription termination, prokaryote ribosome binding site, and restriction enzymes cleavage sites. CAI value^84^ and GC content of the adapted sequence were calculated. Furthermore, XhoI and NdeI restriction sites were introduced to the N and C terminal of the sequence, respectively. Finally, the adapted nucleotide sequence corresponding to the designed vaccine construct was cloned into the *E. coli* pET28a(+) vector by using SnapGene 4.2 tool (https://snapgene.com/) to ensure the vaccine expression.

### Molecular docking between the vaccine and TLR4 receptor

Molecular docking is a computational method which involves the interaction of a ligand molecule to the receptor molecules to give the stable adduct^85^. It can also be used to predict the binding affinity between these two molecules in terms of scoring function. Toll-like receptor 4 (TLR4) is well-known for recognizing ligands from many Gram-negative bacteria. Besides, TLR4 activation may contribute to upregulation of TSPO/MBR related proteins through overproduction of cytokines^69^. Therefore, TLR4 was selected as the receptor and obtained from the RCSB PDB database (PDB ID: 4G8A)^86^ while the vaccine model was used as a ligand. Finally, the binding affinity between the multiepitope vaccine and TLR4 receptor was calculated through molecular docking using the ClusPro 2.0 server^87^. This server completed the task in three successive steps such as rigid body docking, clustering of lowest energy structure, and structural refinement by energy minimization^87^. The best docked complex was selected based on the lowest energy scoring and docking efficiency.

### Dynamic simulations for vaccine stability

Molecular dynamic simulation (MDS) is essential to determine the stability of receptor-ligand complex^35^. The MDS between TLR4 (as receptor) and vaccine (as ligand) was performed using AMBER14 force field^88^ in YASARA Dynamics^89^. Briefly, the complex was cleaned and optimized for hydrogen bond networks before the simulation followed by the creation of cubic simulation cell solvated with TIP3P water (density: 0.997 g/L^−1^). A cut-off radius of 8.0 Å was used for short-range van der Waals and Coulomb interactions at physiological conditions (298 K, pH 7.4, 0.9% NaCl). Timestep of 1.25 fs was used and simulation snapshots were captured at every 100 ps. The MDS was executed for the period of 5 ns until the TLR4-Vaccine complex reaches a stable state. Finally, the trajectories generated from the simulation were analysed for the stability of the complex in terms of the root mean square deviation (RMSD) and root mean square fluctuation (RMSF) for backbone and side chain, respectively. Besides, comparison between initial and final protein backbones for hydrogen bonds was evaluated with BIOVIA Discovery Studio 2017.

### Immune simulation for vaccine efficacy

*In silico* immune simulations were carried out using the C-ImmSim server (http://150.146.2.1/C-IMMSIM/index.php) to assess the immunogenic profile of multiepitope vaccine in real-life^44^. C-ImmSim is an agent-based dynamic simulator for immune responses which uses position-specific scoring matrix (PSSM) and machine learning techniques for the prediction of epitopes and immune interactions, respectively^44^. For most vaccines currently in use, the minimum recommended interval between dose 1 and dose 2 is 4 weeks^90^. Therefore, three injections, containing 1000 vaccine proteins each, were administered (the parameters were set in C-ImmSim immune simulator) four weeks apart at 1, 84 and 168 time-steps (each time-step is equivalent to 8 hours in real-life and time-step 1 is injection at time = 0) with total 1050 simulation steps. All other simulation parameters were kept defaults. Furthermore, 12 injections of the designed peptide were given four weeks apart to simulate repeated exposure to the antigen seen in a typical endemic area so as to probe the clonal selection. The Simpson index, D (a measure of diversity) was interpreted from the plot.

## Data availability

The data with particular importance are embedded within the submitted manuscript and the rests are provided as supplementary files. The sequence of proteins used in this study can be retrieved from IMG/M, RCSB PDB and NCBI database using their corresponding accession numbers.

## Author Contributions

Z.N and U.K.A conceived and designed the study; Z.N and F.A carried out the major immunoinformatic analysis; Z.N and U.K.A wrote the manuscript; Z.N prepared the graphs and illustrations; Z.N, S.M.R.R, S.A.K, S.B.S, M.M.S, and Z.H performed the antigenicity and epitope predictions; M.M.K and M.M.R contributed to the critical revision of the manuscript; U.K.A supervised the whole work; and all authors approved the final manuscript.

