## Supplementary Tables and Figures for "Immunoinformatic and dynamic simulation-based designing of a multi-epitope vaccine against emerging pathogen *Elizabethkingia anophelis*"

**Supplementary Files**

**Supplementary Figure 1**: Molecular docking between T-cell epitopes and MHC alleles. (A) Docking between CTLs and HLA-B*53:01 (1. Control Ligand, 2. FIIFFNFKL, 3. ISLWFIIFF, 4. LISLWFIIF, 5. LWFIIFFNF, and 6. VLISLWFII), (B) Docking between HTLs and HLA-DRB5*01:01 (1. Control Ligand, 2. AFYFIWRAKENRFKH, 3. FIWRAKENRFKHRQK, 4. FYFIWRAKENRFKHR, 5. IAFYFIWRAKENRFK, 6. IWRAKENRFKHRQKA, and 7. WRAKENRFKHRQKAL)

**Supplementary Figure 2:** Immune simulation of repeated antigen exposure using C-ImmSim server.

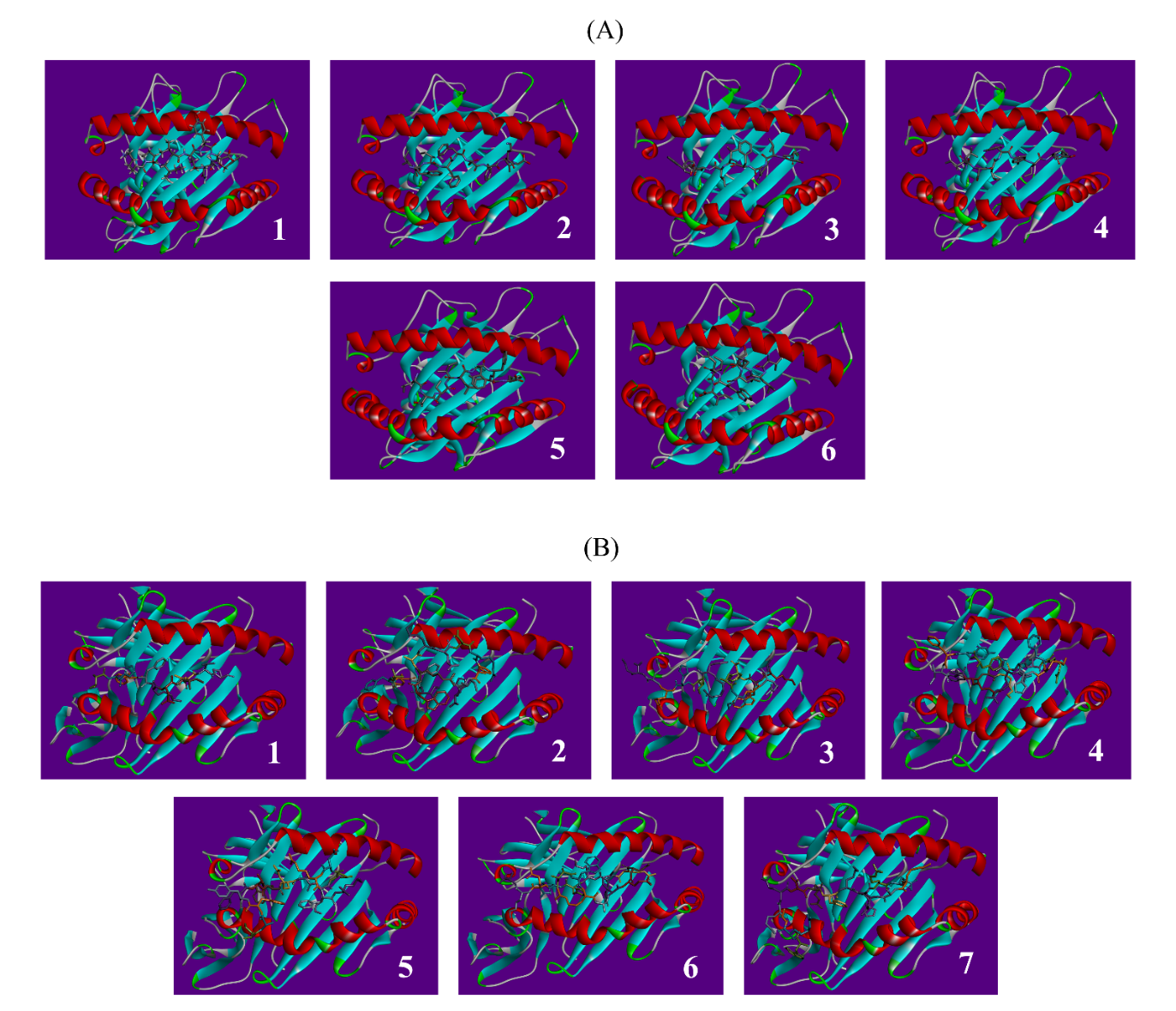

**Supplementary Figure 1: Molecular docking between T-cell epitopes and MHC alleles.** (A) Docking between CTLs and HLA-B*53:01 (1. Control Ligand, 2. FIIFFNFKL, 3. ISLWFIIFF, 4. LISLWFIIF, 5. LWFIIFFNF, and 6. VLISLWFII), (B) Docking between HTLs and HLA-DRB5*01:01 (1. Control Ligand, 2. AFYFIWRAKENRFKH, 3. FIWRAKENRFKHRQK, 4. FYFIWRAKENRFKHR, 5. IAFYFIWRAKENRFK, 6. IWRAKENRFKHRQKA, and 7. WRAKENRFKHRQKAL)

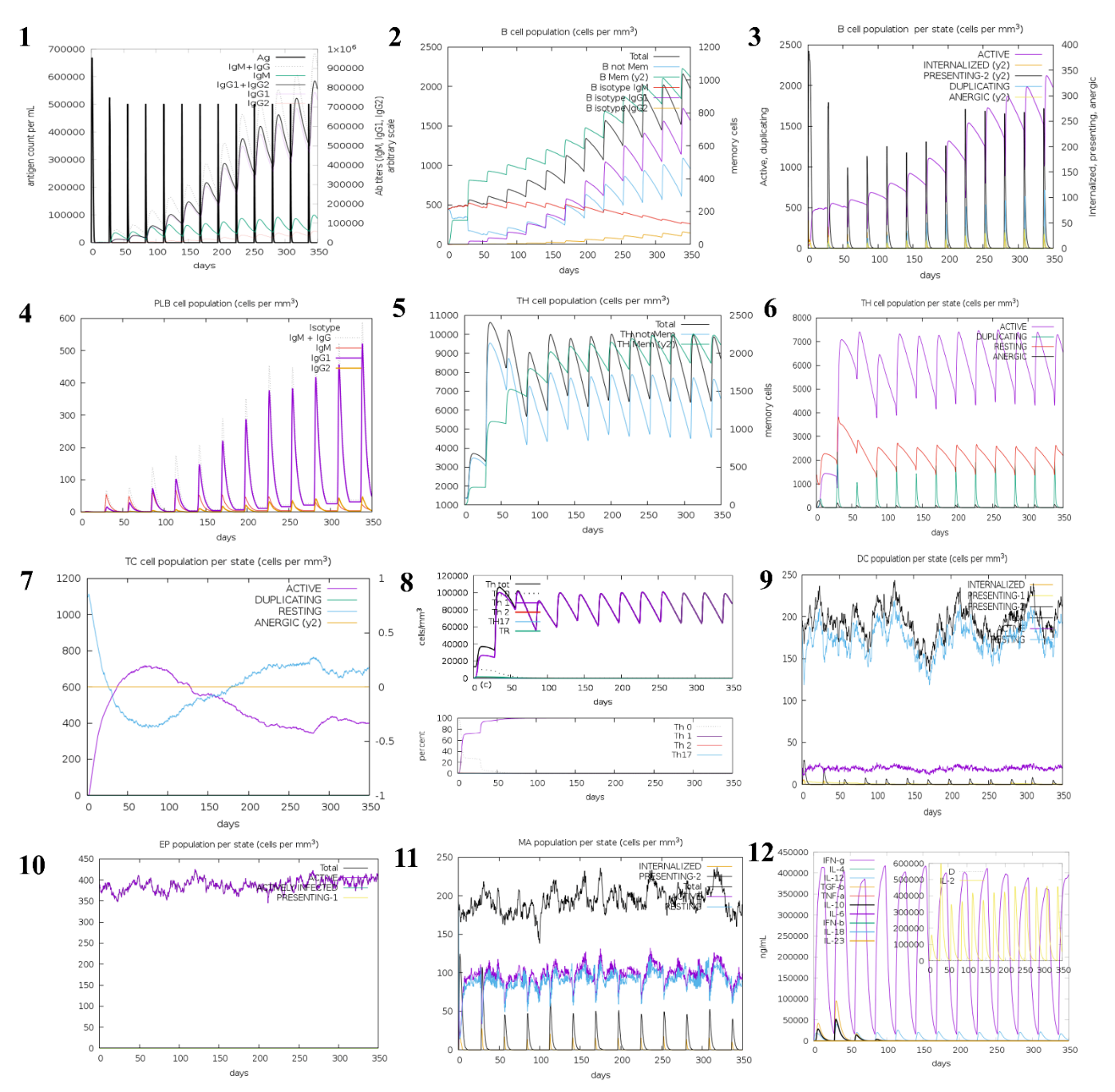

**Supplementary Figure 2:** Immune simulation of repeated antigen exposure using C-ImmSim server.

**Supplementary Tables:**

**Supplementary Table 1:** List of primarily assessed 49 CTL from *E. anophelis* translocator protein.

**Supplementary Table 2:** List of predicted 17 unique HTL epitopes from translocator protein.

**Supplementary Table 3:** List of predicted 4 unique LBL epitopes from EA translocator protein.

**Supplementary Table 4:** Population coverage of T-cell epitopes predicted from translocator protein.

**Supplementary Table 5:** Selected CTL epitopes with their binding alleles (Consensus: PR≤2).

**Supplementary Table 6:** Selected T-cell epitopes with their respective binding alleles.

**Supplementary Table 7:** Tertiary model prediction of the final vaccine construct.

**Supplementary Table 8:** Tertiary structure refinement result of the final vaccine protein.

**Supplementary Table 9:** Disulfide engineering of the final multiepitope vaccine.

**Supplementary Table 10**: Energy scores of the top 10 vaccine-receptor docked complexes.

**Supplementary Table 11**: Molecular dynamic simulations of the vaccine-receptor complex.

**Supplementary Table 1:** List of primarily assessed 49 CTL from *E. anophelis*-derived translocator protein

| **Supertype** | **CTL Epitope** | **Combined** | **Antigenicity** | **Immunogenicity** | **Allergenicity** | **Toxicity** |
| --- | --- | --- | --- | --- | --- | --- |
| A2 | ACISFHVFV | 0.567 | 1.3117 | 0.12442 | No | Non-Toxin |
| A3 | ALAACISFH | 1.0654 | 0.7449 | 0.05363 | No | Non-Toxin |
| B62 | ALTYYYIIF | 0.8121 | 1.9704 | 0.19188 | No | Non-Toxin |
| A2 | ASYLILPTL | 0.6963 | 0.5171 | 0.12012 | No | Non-Toxin |
| B58 | CISFHVFVF | 1.0457 | 2.0933 | 0.25738 | No | Non-Toxin |
| B39 | FHVFVFFII | 1.4583 | 2.8494 | 0.45816 | No | Non-Toxin |
| A2 | FIIFFNFKL | 1.3485 | 3.4177 | 0.24171 | No | Non-Toxin |
| A2 | FLFRTIFLM | 1.3221 | 1.0356 | 0.34548 | No | Non-Toxin |
| A2 | FLMYIFSGI | 1.4001 | 0.9096 | 0.05926 | No | Non-Toxin |
| A2 | FSGIAFYFI | 0.575 | 1.5923 | 0.3585 | No | Non-Toxin |
| A2 | FVFFIIHAL | 1.3246 | 1.5522 | 0.46084 | No | Non-Toxin |
| A2 | FVLISLWFI | 1.2304 | 2.7851 | 0.21412 | No | Non-Toxin |
| A3 | GIAFYFIWR | 1.245 | 2.3643 | 0.47884 | No | Non-Toxin |
| A3 | HVFVFFIIH | 0.7904 | 1.8254 | 0.49382 | No | Non-Toxin |
| A24 | IFSGIAFYF | 1.7223 | 0.9789 | 0.24347 | No | Non-Toxin |
| A24 | IFVLISLWF | 1.7749 | 2.3336 | 0.09617 | No | Non-Toxin |
| A2 | IIFFNFKLL | 0.5147 | 2.3287 | 0.07122 | No | Non-Toxin |
| B58 | IIFVLISLW | 1.3768 | 1.7866 | 0.04792 | No | Non-Toxin |
| A3 | IILFFLFKR | 1.245 | 1.6308 | 0.19056 | No | Non-Toxin |
| A2 | IITWVLTII | 0.6861 | 0.9575 | 0.37577 | No | Non-Toxin |
| A2 | ILFFLFKRV | 0.9363 | 0.8553 | 0.10344 | No | Non-Toxin |
| B58 | ISFHVFVFF | 1.5894 | 2.3259 | 0.32419 | No | Non-Toxin |
| B58 | ISLWFIIFF | 1.5548 | 4.3489 | 0.63929 | No | Non-Toxin |
| A2 | ITWVLTIIL | 0.8073 | 1.2249 | 0.32926 | No | Non-Toxin |
| B62 | LFRTIFLMY | 0.951 | 0.7531 | 0.1837 | No | Non-Toxin |
| B62 | LISLWFIIF | 0.9822 | 3.8608 | 0.45112 | No | Non-Toxin |
| B58 | LTIILFFLF | 1.5784 | 3.6595 | 0.36884 | No | Non-Toxin |
| A2 | LTYYYIIFV | 1.3183 | 1.9068 | 0.29748 | No | Non-Toxin |
| A24 | LWFIIFFNF | 1.802 | 4.9377 | 0.50674 | No | Non-Toxin |
| A3 | MVLAWGFWQ | 0.7373 | 0.916 | 0.51159 | No | Non-Toxin |
| A24 | NFLFRTIFL | 0.9304 | 1.0001 | 0.38186 | No | Non-Toxin |
| A2 | NIITWVLTI | 1.0177 | 1.0001 | 0.35014 | No | Non-Toxin |
| A1 | PANSFNVWY | 1.1007 | 0.6562 | 0.1036 | No | Non-Toxin |
| B27 | RQIDWKALA | 0.9155 | 1.3007 | 0.10476 | No | Non-Toxin |
| A24 | SFHVFVFFI | 1.541 | 2.3929 | 0.3721 | No | Non-Toxin |
| B58 | SGIAFYFIW | 1.8475 | 2.8185 | 0.36965 | No | Non-Toxin |
| B62 | SNFLFRTIF | 0.7026 | 0.676 | 0.30008 | No | Non-Toxin |
| A24 | SYLILPTLL | 1.8158 | 0.7726 | 0.13536 | No | Non-Toxin |
| A3 | TIILFFLFK | 1.5034 | 2.2595 | 0.31528 | No | Non-Toxin |
| A24 | TWVLTIILF | 1.7598 | 1.6572 | 0.27116 | No | Non-Toxin |
| A24 | TYYYIIFVL | 1.7998 | 1.7767 | 0.37288 | No | Non-Toxin |
| A2 | VLAWGFWQL | 1.3194 | 1.0316 | 0.49805 | No | Non-Toxin |
| A2 | VLISLWFII | 1.0013 | 3.0315 | 0.251 | No | Non-Toxin |
| A2 | VLTIILFFL | 1.2793 | 2.4769 | 0.43288 | No | Non-Toxin |
| A24 | WFIIFFNFK | 0.7288 | 3.9403 | 0.46426 | No | Non-Toxin |
| A24 | WVLTIILFF | 1.2366 | 2.3723 | 0.34938 | No | Non-Toxin |
| A26 | YIIFVLISL | 1.4937 | 1.5534 | 0.20642 | No | Non-Toxin |
| A24 | YYIIFVLIS | 0.6485 | 1.5585 | 0.39838 | No | Non-Toxin |
| A24 | YYYIIFVLI | 1.94 | 2.0943 | 0.40088 | No | Non-Toxin |

**Supplementary Table 2:** List of predicted 17 unique HTL epitopes from *E. anophelis*-derived translocator protein

| **Peptide ID** | **HTL Epitope** | **IFN-γ Prediction** | **IL-4 Prediction** | **IL-10 Prediction** |
| --- | --- | --- | --- | --- |
| 3 | AFYFIWRAKENRFKH | Positive | IL4 inducer | IL10 inducer |
| 16 | FFLFKRVSKTASYLI | Positive | IL4 inducer | IL10 inducer |
| 21 | FIWRAKENRFKHRQK | Positive | IL4 inducer | IL10 inducer |
| 25 | FLFKRVSKTASYLIL | Positive | IL4 inducer | IL10 inducer |
| 29 | FSGIAFYFIWRAKEN | Positive | IL4 inducer | IL10 inducer |
| 33 | FYFIWRAKENRFKHR | Positive | IL4 inducer | IL10 inducer |
| 34 | GIAFYFIWRAKENRF | Positive | IL4 inducer | IL10 inducer |
| 37 | IAFYFIWRAKENRFK | Positive | IL4 inducer | IL10 inducer |
| 41 | IFSGIAFYFIWRAKE | Positive | IL4 inducer | IL10 inducer |
| 49 | ILFFLFKRVSKTASY | Positive | IL4 inducer | IL10 inducer |
| 55 | IWRAKENRFKHRQKA | Positive | IL4 inducer | IL10 inducer |
| 58 | KENRFKHRQKALTYY | Positive | IL4 inducer | IL10 inducer |
| 94 | RAKENRFKHRQKALT | Positive | IL4 inducer | IL10 inducer |
| 102 | SGIAFYFIWRAKENR | Positive | IL4 inducer | IL10 inducer |
| 127 | WRAKENRFKHRQKAL | Positive | IL4 inducer | IL10 inducer |
| 130 | YFIWRAKENRFKHRQ | Positive | IL4 inducer | IL10 inducer |
| 131 | YIFSGIAFYFIWRAK | Positive | IL4 inducer | IL10 inducer |

**Supplementary Table 3:** List of predicted 4 unique LBL epitopes from EA translocator protein

| **Linear B-lymphocyte Epitope** | **Residues** | **Start** | **End** | **Score** | **Antigenicity** | **Allergenicity** | **Toxicity** |
| --- | --- | --- | --- | --- | --- | --- | --- |
| MKRQIDWKALAA | 12 | 1 | 12 | 0.794 | 0.8589 | No | No |
| WRAKENRFKHRQK | 13 | 71 | 83 | 0.764 | 1.0623 | No | Yes |
| FNFKLLDL | 8 | 103 | 110 | 0.695 | 2.822 | Yes | No |
| ALTPANSFNVWYNQLDKPLFTPPSNF | 26 | 26 | 51 | 0.686 | 0.4166 | No | No |

**Supplementary Table 4:** Population coverage of T-cell epitopes predicted from EA translocator protein

| **MHC class I** | | **MHC class II** | | **MHC class I and II (combined)** | |
| --- | --- | --- | --- | --- | --- |
| **Regions** | **Coverage** | **Regions** | **Coverage** | **Regions** | **Coverage** |
| American Samoa | 98.99% | Algeria | 76.97% | Algeria | 76.97% |
| American Samoa Polynesian | 98.99% | Algeria Arab | 76.97% | Algeria Arab | 76.97% |
| Argentina | 99.97% | Argentina | 59.02% | American Samoa | 98.99% |
| Argentina Amerindian | 99.97% | Argentina Amerindian | 45.86% | American Samoa Polynesian | 98.99% |
| Australia | 99.01% | Argentina Caucasoid | 73.68% | Argentina | 99.99% |
| Australia Australian Aborigines | 97.91% | Australia | 29.95% | Argentina Amerindian | 99.98% |
| Australia Caucasoid | 99.99% | Australia Australian Aborigines | 29.95% | Argentina Caucasoid | 73.68% |
| Austria | 99.97% | Austria | 75.20% | Australia | 99.30% |
| Austria Caucasoid | 99.97% | Austria Caucasoid | 75.20% | Australia Australian Aborigines | 98.54% |
| Belgium | 99.69% | Belarus | 27.09% | Australia Caucasoid | 99.99% |
| Belgium Caucasoid | 99.69% | Belarus Caucasoid | 27.09% | Austria | 99.99% |
| Brazil | 99.53% | Belgium | 54.84% | Austria Caucasoid | 99.99% |
| Brazil Amerindian | 99.54% | Belgium Caucasoid | 54.84% | Belarus | 27.09% |
| Brazil Caucasoid | 99.53% | Bolivia | 76.86% | Belarus Caucasoid | 27.09% |
| Brazil Mixed | 99.70% | Bolivia Amerindian | 76.86% | Belgium | 99.86% |
| Bulgaria | 99.99% | Borneo | 61.68% | Belgium Caucasoid | 99.86% |
| Bulgaria Caucasoid | 99.96% | Borneo Austronesian | 61.68% | Bolivia | 76.86% |
| Bulgaria Other | 100.00% | Brazil | 60.62% | Bolivia Amerindian | 76.86% |
| Burkina Faso | 84.97% | Brazil Amerindian | 51.91% | Borneo | 61.68% |
| Burkina Faso Black | 84.97% | Brazil Caucasoid | 78.49% | Borneo Austronesian | 61.68% |
| Cameroon | 98.73% | Brazil Mixed | 67.10% | Brazil | 99.81% |
| Cameroon Black | 98.73% | Brazil Mulatto | 68.98% | Brazil Amerindian | 99.78% |
| Cape Verde | 99.67% | Bulgaria | 78.47% | Brazil Caucasoid | 99.90% |
| Cape Verde Black | 99.67% | Bulgaria Caucasoid | 78.47% | Brazil Mixed | 99.90% |
| Central Africa | 97.50% | Cameroon | 63.28% | Brazil Mulatto | 68.98% |
| Central African Republic | 38.72% | Cameroon Black | 63.28% | Bulgaria | 100.00% |
| Central African Republic Black | 38.72% | Canada | 34.15% | Bulgaria Caucasoid | 99.99% |
| Central America | 7.95% | Canada Amerindian | 34.15% | Bulgaria Other | 100.00% |
| Chile | 99.58% | Cape Verde | 73.84% | Burkina Faso | 84.97% |
| Chile Amerindian | 100.00% | Cape Verde Black | 73.84% | Burkina Faso Black | 84.97% |
| Chile Mixed | 97.58% | Central Africa | 63.91% | Cameroon | 99.53% |
| China | 98.51% | Central African Republic | 85.56% | Cameroon Black | 99.53% |
| China Oriental | 98.51% | Central African Republic Black | 85.56% | Canada | 34.15% |
| Colombia | 82.35% | Central America | 49.46% | Canada Amerindian | 34.15% |
| Colombia Amerindian | 83.46% | Chile | 61.75% | Cape Verde | 99.91% |
| Colombia Black | 86.73% | Chile Amerindian | 67.40% | Cape Verde Black | 99.91% |
| Colombia Mestizo | 77.57% | Chile Mixed | 53.91% | Central Africa | 99.10% |
| Croatia | 99.92% | China | 58.80% | Central African Republic | 91.15% |
| Croatia Caucasoid | 99.92% | China Oriental | 58.80% | Central African Republic Black | 91.15% |
| Cuba | 99.40% | Colombia | 52.26% | Central America | 53.48% |
| Cuba Caucasoid | 99.63% | Colombia Amerindian | 52.65% | Chile | 99.84% |
| Cuba Mulatto | 98.98% | Colombia Black | 50.13% | Chile Amerindian | 100.00% |
| Czech Republic | 99.88% | Colombia Mestizo | 45.68% | Chile Mixed | 98.88% |
| Czech Republic Caucasoid | 99.88% | Congo | 65.71% | China | 99.39% |
| East Africa | 97.89% | Congo Black | 65.71% | China Oriental | 99.39% |
| East Asia | 99.95% | Cook Islands | 73.31% | Colombia | 91.32% |
| Ecuador | 99.78% | Cook Islands Polynesian | 73.31% | Colombia Amerindian | 91.75% |
| Ecuador Amerindian | 99.78% | Costa Rica | 29.44% | Colombia Black | 93.38% |
| England | 99.99% | Costa Rica Mestizo | 29.44% | Colombia Mestizo | 87.82% |
| England Caucasoid | 100.00% | Croatia | 73.83% | Congo | 65.71% |
| England Jew | 30.61% | Croatia Caucasoid | 73.83% | Congo Black | 65.71% |
| Equatorial Guinea | 1.00% | Cuba | 73.68% | Cook Islands | 73.31% |
| Equatorial Guinea Black | 1.00% | Cuba Mixed | 73.68% | Cook Islands Polynesian | 73.31% |
| Europe | 99.99% | Czech Republic | 81.59% | Costa Rica | 29.44% |
| Finland | 100.00% | Czech Republic Caucasoid | 83.23% | Costa Rica Mestizo | 29.44% |
| Finland Caucasoid | 100.00% | Czech Republic Other | 68.21% | Croatia | 99.98% |
| France | 99.99% | Denmark | 56.70% | Croatia Caucasoid | 99.98% |
| France Caucasoid | 99.99% | Denmark Caucasoid | 56.70% | Cuba | 99.84% |
| Georgia | 99.86% | East Africa | 69.97% | Cuba Caucasoid | 99.63% |
| Georgia Caucasoid | 99.94% | East Asia | 66.96% | Cuba Mixed | 73.68% |
| Georgia Kurd | 99.59% | Ecuador | 61.96% | Cuba Mulatto | 98.98% |
| Germany | 100.00% | Ecuador Amerindian | 61.96% | Czech Republic | 99.98% |
| Germany Caucasoid | 100.00% | England | 70.60% | Czech Republic Caucasoid | 99.98% |
| Guatemala | 7.95% | England Caucasoid | 70.60% | Czech Republic Other | 68.21% |
| Guatemala Amerindian | 7.95% | Equatorial Guinea | 78.19% | Denmark | 56.70% |
| Guinea-Bissau | 98.57% | Equatorial Guinea Black | 78.19% | Denmark Caucasoid | 56.70% |
| Guinea-Bissau Black | 98.57% | Ethiopia | 57.14% | East Africa | 99.37% |
| Hong Kong | 97.96% | Ethiopia Black | 57.14% | East Asia | 99.98% |
| Hong Kong Oriental | 97.96% | Europe | 73.60% | Ecuador | 99.92% |
| India | 99.01% | Fiji | 89.37% | Ecuador Amerindian | 99.92% |
| India Asian | 99.01% | Fiji Melanesian | 89.37% | England | 100.00% |
| Indonesia | 90.45% | Finland | 36.96% | England Caucasoid | 100.00% |
| Indonesia Austronesian | 90.45% | Finland Caucasoid | 36.96% | England Jew | 30.61% |
| Iran | 99.51% | France | 71.78% | Equatorial Guinea | 78.41% |
| Iran Persian | 99.51% | France Caucasoid | 71.78% | Equatorial Guinea Black | 78.41% |
| Ireland Northern | 100.00% | Gabon | 51.84% | Ethiopia | 57.14% |
| Ireland Northern Caucasoid | 100.00% | Gabon Black | 51.84% | Ethiopia Black | 57.14% |
| Ireland South | 100.00% | Georgia | 82.30% | Europe | 100.00% |
| Ireland South Caucasoid | 100.00% | Georgia Caucasoid | 82.30% | Fiji | 89.37% |
| Israel | 98.00% | Germany | 77.32% | Fiji Melanesian | 89.37% |
| Israel Arab | 99.42% | Germany Caucasoid | 77.32% | Finland | 100.00% |
| Israel Jew | 97.97% | Greece | 74.21% | Finland Caucasoid | 100.00% |
| Italy | 99.90% | Greece Caucasoid | 74.21% | France | 100.00% |
| Italy Caucasoid | 99.90% | Guatemala | 47.00% | France Caucasoid | 100.00% |
| Ivory Coast | 73.89% | Guatemala Amerindian | 47.00% | Gabon | 51.84% |
| Ivory Coast Black | 73.89% | Guinea-Bissau | 83.92% | Gabon Black | 51.84% |
| Japan | 99.98% | Guinea-Bissau Black | 83.92% | Georgia | 99.98% |
| Japan Oriental | 99.98% | India | 74.93% | Georgia Caucasoid | 99.99% |
| Jordan | 95.31% | India Asian | 74.93% | Georgia Kurd | 99.59% |
| Jordan Arab | 95.31% | Indonesia | 71.27% | Germany | 100.00% |
| Kenya | 98.25% | Indonesia Austronesian | 71.27% | Germany Caucasoid | 100.00% |
| Kenya Black | 98.25% | Iran | 64.46% | Greece | 74.21% |
| Korea; South | 99.94% | Iran Kurd | 68.08% | Greece Caucasoid | 74.21% |
| Korea; South Oriental | 99.94% | Iran Persian | 64.24% | Guatemala | 51.22% |
| Lebanon | 73.79% | Ireland Northern | 70.92% | Guatemala Amerindian | 51.22% |
| Lebanon Mixed | 73.79% | Ireland Northern Caucasoid | 70.92% | Guinea-Bissau | 99.77% |
| Macedonia | 38.74% | Ireland South | 68.08% | Guinea-Bissau Black | 99.77% |
| Macedonia Caucasoid | 38.74% | Ireland South Caucasoid | 68.08% | Hong Kong | 97.96% |
| Malaysia | 84.70% | Israel | 76.95% | Hong Kong Oriental | 97.96% |
| Malaysia Austronesian | 75.79% | Israel Arab | 72.63% | India | 99.75% |
| Malaysia Oriental | 89.96% | Israel Jew | 78.50% | India Asian | 99.75% |
| Mali | 99.24% | Italy | 34.36% | Indonesia | 97.26% |
| Mali Black | 99.24% | Italy Caucasoid | 34.36% | Indonesia Austronesian | 97.26% |
| Martinique | 27.75% | Jamaica | 18.10% | Iran | 99.83% |
| Martinique Black | 27.75% | Jamaica Black | 18.10% | Iran Kurd | 68.08% |
| Mexico | 99.96% | Japan | 63.15% | Iran Persian | 99.83% |
| Mexico Amerindian | 100.00% | Japan Oriental | 63.15% | Ireland Northern | 100.00% |
| Mexico Mestizo | 99.69% | Jordan | 33.82% | Ireland Northern Caucasoid | 100.00% |
| Mongolia | 96.13% | Jordan Arab | 33.82% | Ireland South | 100.00% |
| Mongolia Oriental | 96.13% | Kiribati | 30.94% | Ireland South Caucasoid | 100.00% |
| Morocco | 99.76% | Kiribati Micronesian | 30.94% | Israel | 99.54% |
| Morocco Arab | 99.73% | Korea; South | 63.48% | Israel Arab | 99.84% |
| Morocco Caucasoid | 99.81% | Korea; South Oriental | 63.48% | Israel Jew | 99.56% |
| New Caledonia | 99.27% | Lebanon | 77.88% | Italy | 99.93% |
| New Caledonia Melanesian | 99.27% | Lebanon Arab | 77.88% | Italy Caucasoid | 99.93% |
| North Africa | 99.24% | Macedonia | 76.42% | Ivory Coast | 73.89% |
| North America | 99.94% | Macedonia Caucasoid | 76.42% | Ivory Coast Black | 73.89% |
| Northeast Asia | 98.54% | Malaysia | 65.58% | Jamaica | 18.10% |
| Oceania | 98.72% | Malaysia Austronesian | 65.02% | Jamaica Black | 18.10% |
| Oman | 99.67% | Malaysia Oriental | 72.08% | Japan | 99.99% |
| Oman Arab | 99.67% | Martinique | 71.49% | Japan Oriental | 99.99% |
| Pakistan | 98.68% | Martinique Black | 71.49% | Jordan | 96.89% |
| Pakistan Asian | 98.46% | Mexico | 53.92% | Jordan Arab | 96.89% |
| Pakistan Mixed | 99.07% | Mexico Amerindian | 47.18% | Kenya | 98.25% |
| Papua New Guinea | 99.57% | Mexico Mestizo | 62.79% | Kenya Black | 98.25% |
| Papua New Guinea Melanesian | 99.57% | Mongolia | 69.80% | Kiribati | 30.94% |
| Peru | 100.00% | Mongolia Oriental | 69.80% | Kiribati Micronesian | 30.94% |
| Peru Amerindian | 100.00% | Morocco | 69.15% | Korea; South | 99.98% |
| Peru Mestizo | 1.99% | Morocco Arab | 67.57% | Korea; South Oriental | 99.98% |
| Philippines | 99.36% | Morocco Caucasoid | 74.40% | Lebanon | 94.20% |
| Philippines Austronesian | 99.36% | Nauru | 65.26% | Lebanon Arab | 77.88% |
| Poland | 99.99% | Nauru Micronesian | 65.26% | Lebanon Mixed | 73.79% |
| Poland Caucasoid | 99.99% | Netherlands | 73.95% | Macedonia | 85.55% |
| Portugal | 99.79% | Netherlands Caucasoid | 73.95% | Macedonia Caucasoid | 85.55% |
| Portugal Caucasoid | 99.79% | New Caledonia | 86.29% | Malaysia | 94.73% |
| Romania | 99.96% | New Caledonia Melanesian | 86.29% | Malaysia Austronesian | 91.53% |
| Romania Caucasoid | 99.96% | New Zealand | 76.98% | Malaysia Oriental | 97.20% |
| Russia | 99.88% | New Zealand Polynesian | 76.98% | Mali | 99.24% |
| Russia Caucasoid | 68.80% | Niue | 79.75% | Mali Black | 99.24% |
| Russia Mixed | 66.00% | Niue Polynesian | 79.75% | Martinique | 79.40% |
| Russia Other | 100.00% | North Africa | 72.74% | Martinique Black | 79.40% |
| Russia Siberian | 99.88% | North America | 70.82% | Mexico | 99.98% |
| Rwanda | 25.84% | Northeast Asia | 58.80% | Mexico Amerindian | 100.00% |
| Rwanda Black | 25.84% | Norway | 73.99% | Mexico Mestizo | 99.89% |
| Sao Tome and Principe | 98.47% | Norway Caucasoid | 73.99% | Mongolia | 98.83% |
| Sao Tome and Principe Black | 98.47% | Oceania | 63.12% | Mongolia Oriental | 98.83% |
| Saudi Arabia | 99.66% | Pakistan | 2.19% | Morocco | 99.93% |
| Saudi Arabia Arab | 99.66% | Pakistan Asian | 2.19% | Morocco Arab | 99.91% |
| Scotland | 56.04% | Papua New Guinea | 74.63% | Morocco Caucasoid | 99.95% |
| Scotland Caucasoid | 56.04% | Papua New Guinea Melanesian | 74.63% | Nauru | 65.26% |
| Senegal | 98.70% | Paraguay | 4.90% | Nauru Micronesian | 65.26% |
| Senegal Black | 98.70% | Paraguay Amerindian | 4.90% | Netherlands | 73.95% |
| Serbia | 80.90% | Peru | 60.25% | Netherlands Caucasoid | 73.95% |
| Serbia Caucasoid | 80.90% | Peru Amerindian | 60.25% | New Caledonia | 99.90% |
| Singapore | 98.02% | Philippines | 75.60% | New Caledonia Melanesian | 99.90% |
| Singapore Austronesian | 97.01% | Philippines Austronesian | 75.60% | New Zealand | 76.98% |
| Singapore Oriental | 98.69% | Poland | 76.52% | New Zealand Polynesian | 76.98% |
| South Africa | 99.34% | Poland Caucasoid | 76.52% | Niue | 79.75% |
| South Africa Black | 96.93% | Portugal | 75.75% | Niue Polynesian | 79.75% |
| South Africa Other | 99.90% | Portugal Caucasoid | 75.75% | North Africa | 99.79% |
| South America | 99.82% | Russia | 71.80% | North America | 99.98% |
| South Asia | 99.45% | Russia Caucasoid | 75.84% | Northeast Asia | 99.40% |
| Southeast Asia | 99.74% | Russia Other | 80.01% | Norway | 73.99% |
| Southwest Asia | 97.90% | Russia Siberian | 72.55% | Norway Caucasoid | 73.99% |
| Spain | 99.46% | Rwanda | 52.25% | Oceania | 99.53% |
| Spain Caucasoid | 99.46% | Rwanda Black | 52.25% | Oman | 99.67% |
| Sri Lanka | 63.16% | Samoa | 75.35% | Oman Arab | 99.67% |
| Sri Lanka Asian | 63.16% | Samoa Polynesian | 75.35% | Pakistan | 98.71% |
| Sudan | 98.83% | Sao Tome and Principe | 60.80% | Pakistan Asian | 98.50% |
| Sudan Arab | 77.35% | Sao Tome and Principe Black | 60.80% | Pakistan Mixed | 99.07% |
| Sudan Black | 2.19% | Saudi Arabia | 74.85% | Papua New Guinea | 99.89% |
| Sudan Mixed | 98.97% | Saudi Arabia Arab | 74.85% | Papua New Guinea Melanesian | 99.89% |
| Sweden | 99.98% | Scotland | 69.20% | Paraguay | 4.90% |
| Sweden Caucasoid | 99.98% | Scotland Caucasoid | 69.20% | Paraguay Amerindian | 4.90% |
| Switzerland | 82.53% | Senegal | 79.12% | Peru | 100.00% |
| Switzerland Caucasoid | 82.53% | Senegal Black | 79.12% | Peru Amerindian | 100.00% |
| Taiwan | 99.33% | Singapore | 71.91% | Peru Mestizo | 1.99% |
| Taiwan Oriental | 99.33% | Singapore Austronesian | 71.91% | Philippines | 99.84% |
| Thailand | 99.75% | Slovakia | 0.00% | Philippines Austronesian | 99.84% |
| Thailand Oriental | 99.75% | Slovakia Caucasoid | 0.00% | Poland | 100.00% |
| Tunisia | 99.69% | Slovenia | 73.35% | Poland Caucasoid | 100.00% |
| Tunisia Arab | 99.69% | Slovenia Caucasoid | 73.35% | Portugal | 99.95% |
| Turkey | 69.45% | South Africa | 25.52% | Portugal Caucasoid | 99.95% |
| Turkey Caucasoid | 69.45% | South Africa Black | 25.52% | Romania | 99.96% |
| Uganda | 99.14% | South America | 56.97% | Romania Caucasoid | 99.96% |
| Uganda Black | 99.14% | South Asia | 75.46% | Russia | 99.97% |
| United Arab Emirates | 16.09% | Southeast Asia | 58.44% | Russia Caucasoid | 92.46% |
| United Arab Emirates Arab | 16.09% | Southwest Asia | 50.43% | Russia Mixed | 66.00% |
| United Kingdom | 84.16% | Spain | 75.92% | Russia Other | 100.00% |
| United Kingdom Caucasoid | 84.16% | Spain Caucasoid | 75.84% | Russia Siberian | 99.97% |
| United States | 99.94% | Spain Other | 25.61% | Rwanda | 64.59% |
| United States Amerindian | 99.97% | Sudan | 66.59% | Rwanda Black | 64.59% |
| United States Asian | 99.55% | Sudan Mixed | 66.59% | Samoa | 75.35% |
| United States Black | 99.80% | Sweden | 83.90% | Samoa Polynesian | 75.35% |
| United States Caucasoid | 99.98% | Sweden Caucasoid | 83.90% | Sao Tome and Principe | 99.40% |
| United States Hispanic | 99.90% | Taiwan | 56.66% | Sao Tome and Principe Black | 99.40% |
| United States Mestizo | 99.93% | Taiwan Oriental | 56.66% | Saudi Arabia | 99.92% |
| United States Polynesian | 100.00% | Thailand | 71.58% | Saudi Arabia Arab | 99.92% |
| Venezuela | 98.93% | Thailand Oriental | 71.58% | Scotland | 86.46% |
| Venezuela Amerindian | 98.89% | Tokelau | 57.75% | Scotland Caucasoid | 86.46% |
| Venezuela Caucasoid | 14.73% | Tokelau Polynesian | 57.75% | Senegal | 99.73% |
| Venezuela Mestizo | 13.40% | Tonga | 78.84% | Senegal Black | 99.73% |
| Vietnam | 98.58% | Tonga Polynesian | 78.84% | Serbia | 80.90% |
| Vietnam Oriental | 98.58% | Tunisia | 74.87% | Serbia Caucasoid | 80.90% |
| Wales | 2.97% | Tunisia Arab | 73.79% | Singapore | 99.44% |
| Wales Caucasoid | 2.97% | Tunisia Berber | 83.86% | Singapore Austronesian | 99.16% |
| West Africa | 99.13% | Turkey | 73.58% | Singapore Oriental | 98.69% |
| West Indies | 99.48% | Turkey Caucasoid | 73.58% | Slovakia | 0.00% |
| World | 99.92% | Ukraine | 33.58% | Slovakia Caucasoid | 0.00% |
| Zambia | 98.22% | Ukraine Caucasoid | 33.58% | Slovenia | 73.35% |
| Zambia Black | 98.22% | United States | 71.00% | Slovenia Caucasoid | 73.35% |
| Zimbabwe | 98.32% | United States Amerindian | 34.90% | South Africa | 99.51% |
| Zimbabwe Black | 98.32% | United States Asian | 73.12% | South Africa Black | 97.72% |
|  |  | United States Austronesian | 84.29% | South Africa Other | 99.90% |
|  |  | United States Black | 71.08% | South America | 99.92% |
|  |  | United States Caucasoid | 71.74% | South Asia | 99.86% |
|  |  | United States Hispanic | 69.54% | Southeast Asia | 99.89% |
|  |  | United States Mestizo | 66.84% | Southwest Asia | 98.96% |
|  |  | United States Polynesian | 77.70% | Spain | 99.87% |
|  |  | Venezuela | 3.01% | Spain Caucasoid | 99.87% |
|  |  | Venezuela Mixed | 3.17% | Spain Other | 25.61% |
|  |  | Vietnam | 52.41% | Sri Lanka | 63.16% |
|  |  | Vietnam Oriental | 52.41% | Sri Lanka Asian | 63.16% |
|  |  | West Africa | 78.12% | Sudan | 99.61% |
|  |  | West Indies | 65.46% | Sudan Arab | 77.35% |
|  |  | World | 68.89% | Sudan Black | 2.19% |
|  |  | Zimbabwe | 69.97% | Sudan Mixed | 99.66% |
|  |  | Zimbabwe Black | 69.97% | Sweden | 100.00% |
|  |  |  |  | Sweden Caucasoid | 100.00% |
|  |  |  |  | Switzerland | 82.53% |
|  |  |  |  | Switzerland Caucasoid | 82.53% |
|  |  |  |  | Taiwan | 99.71% |
|  |  |  |  | Taiwan Oriental | 99.71% |
|  |  |  |  | Thailand | 99.93% |
|  |  |  |  | Thailand Oriental | 99.93% |
|  |  |  |  | Tokelau | 57.75% |
|  |  |  |  | Tokelau Polynesian | 57.75% |
|  |  |  |  | Tonga | 78.84% |
|  |  |  |  | Tonga Polynesian | 78.84% |
|  |  |  |  | Tunisia | 99.92% |
|  |  |  |  | Tunisia Arab | 99.92% |
|  |  |  |  | Tunisia Berber | 83.86% |
|  |  |  |  | Turkey | 91.93% |
|  |  |  |  | Turkey Caucasoid | 91.93% |
|  |  |  |  | Uganda | 99.14% |
|  |  |  |  | Uganda Black | 99.14% |
|  |  |  |  | Ukraine | 33.58% |
|  |  |  |  | Ukraine Caucasoid | 33.58% |
|  |  |  |  | United Arab Emirates | 16.09% |
|  |  |  |  | United Arab Emirates Arab | 16.09% |
|  |  |  |  | United Kingdom | 84.16% |
|  |  |  |  | United Kingdom Caucasoid | 84.16% |
|  |  |  |  | United States | 99.98% |
|  |  |  |  | United States Amerindian | 99.98% |
|  |  |  |  | United States Asian | 99.88% |
|  |  |  |  | United States Austronesian | 84.29% |
|  |  |  |  | United States Black | 99.94% |
|  |  |  |  | United States Caucasoid | 99.99% |
|  |  |  |  | United States Hispanic | 99.97% |
|  |  |  |  | United States Mestizo | 99.98% |
|  |  |  |  | United States Polynesian | 100.00% |
|  |  |  |  | Venezuela | 98.96% |
|  |  |  |  | Venezuela Amerindian | 98.89% |
|  |  |  |  | Venezuela Caucasoid | 14.73% |
|  |  |  |  | Venezuela Mestizo | 13.40% |
|  |  |  |  | Venezuela Mixed | 3.17% |
|  |  |  |  | Vietnam | 99.32% |
|  |  |  |  | Vietnam Oriental | 99.32% |
|  |  |  |  | Wales | 2.97% |
|  |  |  |  | Wales Caucasoid | 2.97% |
|  |  |  |  | West Africa | 99.81% |
|  |  |  |  | West Indies | 99.82% |
|  |  |  |  | World | 99.97% |
|  |  |  |  | Zambia | 98.22% |
|  |  |  |  | Zambia Black | 98.22% |
|  |  |  |  | Zimbabwe | 99.49% |
|  |  |  |  | Zimbabwe Black | 99.49% |

**Supplementary Table 5:** Selected CTL epitopes with their respective binding alleles (Consensus: PR≤2).

| **Cytotoxic T lymphocyte** | **Epitopes** | **MHC Alleles** |
| --- | --- | --- |
|  | FIIFFNFKL | HLA-A*02:17, HLA-A*02:02, HLA-A*02:01, HLA-A*69:01, HLA-A*02:50, HLA-A*02:06, HLA-A*68:02, HLA-A*25:01, HLA-A*02:12, HLA-B*35:03, HLA-E*01:03, HLA-B*15:09, HLA-B*15:02, HLA-A*02:11, **HLA-B*53:01**, HLA-A*32:01, HLA-B*39:01, HLA-A*26:01, HLA-A*02:19, HLA-A*26:03, HLA-A*02:03, HLA-A*02:16, HLA-B*42:01, HLA-A*29:02, HLA-A*68:01, HLA-A*23:01, HLA-A*26:02, HLA-B*08:02 |
|  | ISLWFIIFF | HLA-B*58:01, HLA-B*15:17, HLA-B*58:02, HLA-B*57:01, **HLA-B*53:01**, HLA-A*23:01, HLA-A*24:02, HLA-B*15:03, HLA-A*32:01, HLA-B*35:01, HLA-A*69:01, HLA-A*02:06, HLA-A*24:03, HLA-B*18:01, HLA-A*01:01 |
|  | LTIILFFLF | HLA-B*15:17, HLA-A*23:01, HLA-B*58:01, HLA-A*24:02, HLA-A*29:02, HLA-B*57:01, HLA-A*32:01, HLA-B*15:01, HLA-A*26:01, HLA-A*80:01, HLA-A*26:02, HLA-A*32:15, HLA-A*01:01, HLA-A*24:03, HLA-B*35:01, HLA-A*02:06, **HLA-B*53:01**, HLA-B*51:01, HLA-A*66:01, HLA-A*68:01, HLA-A*26:03, HLA-B*58:02, HLA-E*01:03, HLA-B*18:01, HLA-A*69:01, HLA-A*25:01, HLA-A*68:02 |
|  | VLISLWFII | HLA-A*02:01, HLA-A*24:02, HLA-A*02:12, HLA-A*02:16, HLA-A*23:01, HLA-A*02:06, HLA-A*02:19, HLA-A*02:11, HLA-A*02:02, HLA-E*01:03, HLA-A*02:03, HLA-B*51:01, HLA-A*32:01, HLA-A*02:17, **HLA-B*53:01** |
|  | LISLWFIIF | HLA-A*32:15, HLA-B*15:01, HLA-B*15:03, HLA-B*35:01, **HLA-B*53:01**, HLA-A*24:02, HLA-A*32:01, HLA-A*23:01, HLA-A*01:01 |
|  | LWFIIFFNF | HLA-A*23:01, HLA-A*24:02, HLA-A*24:03, HLA-A*32:15, HLA-A*29:02, HLA-A*32:01, HLA-B*15:03, HLA-B*35:01, **HLA-B*53:01**, HLA-A*33:01, HLA-B*18:01, HLA-B*51:01, HLA-E*01:03, HLA-B*15:01, HLA-B*08:02, HLA-C*04:01, HLA-B*40:02 |
| **Helper T lymphocyte** | AFYFIWRAKENRFKH | HLA-DRB1*13:21, HLA-DRB1*08:01, HLA-DRB1*04:02, HLA-DRB1*13:23, HLA-DRB1*11:14, HLA-DRB1*13:07, HLA-DRB1*11:20, HLA-DRB1*08:17, HLA-DRB1*11:28, HLA-DRB1*13:05, HLA-DRB1*08:13, HLA-DRB1*08:06, HLA-DRB1*08:02, HLA-DRB5*01:05, **HLA-DRB5*01:01**, HLA-DRB1*03:09 |
|  | FIWRAKENRFKHRQK | HLA-DRB1*04:02, HLA-DRB1*13:23, HLA-DRB1*11:14, HLA-DRB1*11:20, HLA-DRB1*08:13, HLA-DRB5*01:05, **HLA-DRB5*01:01** |
|  | FYFIWRAKENRFKHR | HLA-DRB1*13:21, HLA-DRB1*08:01, HLA-DRB1*04:02, HLA-DRB1*13:23, HLA-DRB1*11:14, HLA-DRB1*13:07, HLA-DRB1*11:20, HLA-DRB1*08:17, HLA-DRB1*11:28, HLA-DRB1*13:05, HLA-DRB1*08:13, HLA-DRB1*08:02, HLA-DRB1*08:06, HLA-DRB5*01:05, **HLA-DRB5*01:01**, HLA-DRB1*03:09 |
|  | IAFYFIWRAKENRFK | HLA-DRB1*13:21, HLA-DRB1*08:01, HLA-DRB1*04:02, HLA-DRB1*13:23, HLA-DRB1*11:14, HLA-DRB1*13:07, HLA-DRB1*11:20, HLA-DRB1*08:17, HLA-DRB1*11:28, HLA-DRB1*13:05, HLA-DRB1*08:02, HLA-DRB1*08:13, HLA-DRB1*15:06, HLA-DRB1*08:04, HLA-DRB1*08:06, HLA-DRB5*01:05, **HLA-DRB5*01:01**, HLA-DRB1*11:21, HLA-DRB1*11:02, HLA-DRB1*13:22, HLA-DRB1*03:09 |
|  | IWRAKENRFKHRQKA | HLA-DRB1*04:02, HLA-DRB1*13:23, HLA-DRB1*11:14, HLA-DRB1*11:20, HLA-DRB5*01:05, **HLA-DRB5*01:01**, HLA-DRB1*08:13 |
|  | YFIWRAKENRFKHRQ | HLA-DRB1*04:02, HLA-DRB1*13:23, HLA-DRB1*11:14, HLA-DRB1*11:20, HLA-DRB1*08:13, HLA-DRB5*01:05, **HLA-DRB5*01:01**, HLA-DRB1*08:02 |
|  | WRAKENRFKHRQKAL | HLA-DRB1*04:02, HLA-DRB1*13:23, HLA-DRB1*11:14, HLA-DRB1*11:20, HLA-DRB5*01:05, **HLA-DRB5*01:01**, HLA-DRB1*08:13 |

**Supplementary Table 6:** Selected T-cell epitopes with their respective binding alleles

| **Ligands** | **Binding Affinity**  **(Kcal/mol)** | **Hydrogen Bonds** | **Interacting**  **Residues** |
| --- | --- | --- | --- |
| **CTL Control** | **-8.5** | 10 | Lys1, Ile3, Thr143, Gln155, Tyr159, Tyr171, Tyr99, Phe9, Gln5 |
| FIIFFNFKL | -8.6 | 6 | Leu9, Thr73, A:Tyr99, Tyr159, Phe4 |
| ISLWFIIFF | -9.3 | 10 | Tyr84, Arg97, Lys146, Trp147, Ser2, Ile6, Leu3 |
| LISLWFIIF | -10.0 | 22 | Leu1, Ile2, Ile7, Phe9, Asn63, Asn70, Asn77, Arg97, Tyr99, Thr143, Tyr123, Tyr9, Tyr171, Trp5, Ser3 |
| LTIILFFLF | -8.1 | 10 | Phe9, Asn70, Thr73, Trp147, Tyr123, Thr143, Leu5, Leu8 |
| LWFIIFFNF | -7.3 | 9 | Leu1, Phe9, Asn70, Gln155, Ile4, Tyr123, Tyr84, Ile5, Phe3 |
| VLISLWFII | -7.1 | 16 | Trp6, Tyr9, Arg62, Asn70, Arg97, Tyr99, Trp147, Asn63, Leu5, Phe7, Ile9, Ser4, Leu5, Val1 |
| **HTL Control** | 8.5 | 12 | Asn62, Tyr213, Trp261, Arg271, Thr277, Val413, Pro415, Ile412, Thr414, Asn411 |
| AFYFIWRAKENRFKH | 7.2 | 12 | Ala1, Trp6, Arg12, His15, Asn62, Arg271, Thr277, Phe13, Asp270 |
| FIWRAKENRFKHRQK | 6.3 | 19 | Trp3, Asn8, Arg13, Gln9, Arg50, Ser53, Arg50, Asp270, Glu7, Gly58, Asn62, Phe10, Gly284, Glu7, Lys15, His12, Lys6 |
| FYFIWRAKENRFKHR | 6.9 | 14 | Trp5, Asn10, Arg15, Asn62, Arg271, Asn282, Trp5, Asn62, Asn69, Arg271, Ser53, Glu11, Thr277, Lys8, Arg6 |
| IAFYFIWRAKENRFK | 7.3 | 11 | Phe14, Gln9, Asn69, Arg271, Asn282, Ala9, Glu11, Lys10, Trp7 |
| IWRAKENRFKHRQKA | 6.5 | 18 | Arg12, Gln13, Lys14, Ala15, Gln9, Ser53, Tyr213, Arg271, Asn282, Gly284, Arg50, Lys10, Arg8, Glu6 |
| YFIWRAKENRFKHRQ | 7.3 | 24 | Gln9, Ser53, Glu55, Asn62, Tyr213, His281, Asn282, Arg5, Asn9, Arg14, Gln15, Glu8, His13, Phe2, Ala6, Phe51, Asp270 |
| WRAKENRFKHRQKAL | 7.0 | 26 | Trp1, Arg7, Phe8, Leu15, Ser53, Asn62, Asn69, Tyr213, Arg271, His281, Asn282, Thr221, Asp270, Glu5, His10, Gln12, Arg11, Ala14 |

**Supplementary Table 7:** Tertiary model prediction of the final vaccine construct

| **Parameters** | **Results** |
| --- | --- |
| Number of predicted domains | 3 |
| Best template | 1dd3A, p-value 8.67e-04 |
| Overall uGDT (GDT) | 178 (46) |
| Residues modeled | 384(100%) |
| Positions predicted as disordered | 56(14%) |
| Secondary structure | 43%H, 11%E, 44%C |
| Solvent access | 50%E, 28%M, 21%B |

**Supplementary Table 8:** Tertiary structure refinement result of the final vaccine protein

| **Model** | **GDT-HA** | **RMSD** | **MolProbity** | **Clash Score** | **Poor Rotamers** | **Rama Favored** |
| --- | --- | --- | --- | --- | --- | --- |
| Initial | 1.0000 | 0.000 | 3.568 | 133.8 | 4.7 | 92.4 |
| MODEL 1 | 0.9121 | 0.557 | 2.256 | 23.8 | 0.7 | 94.2 |
| MODEL 2 | 0.9128 | 0.554 | 2.316 | 21.7 | 1.3 | 94.2 |
| MODEL 3 | 0.9206 | 0.527 | 2.502 | 23.9 | 2.0 | 94.0 |
| MODEL 4 | 0.9089 | 0.565 | 2.221 | 21.7 | 1.0 | 94.2 |
| MODEL 5 | 0.9076 | 0.571 | 2.470 | 24.5 | 2.0 | 94.8 |

**Supplementary Table 9:** Disulfide engineering of the final multiepitope vaccine

| **Res1 Seq#** | **Res1 AA** | **Res2 Seq#** | **Res2 AA** | **Chi3** | **Energy** | **B-Factors** |
| --- | --- | --- | --- | --- | --- | --- |
| 5 | SER | 8 | GLU | 120.57 | 4 | 0 |
| 16 | MET | 20 | GLU | 112.57 | 2.59 | 0 |
| 20 | GLU | 168 | ARG | 106.05 | 6.98 | 0 |
| 32 | PHE | 36 | ALA | 87.96 | 4.31 | 0 |
| 41 | ALA | 222 | GLU | 121.38 | 2.85 | 0 |
| 54 | VAL | 134 | ALA | 106.92 | 4.23 | 0 |
| 55 | GLU | 133 | ALA | 88.56 | 2.28 | 0 |
| 60 | GLN | 136 | ALA | -73.74 | 3.74 | 0 |
| 61 | SER | 135 | LYS | -59.3 | 4.44 | 0 |
| 61 | SER | 164 | ALA | -105.43 | 3.65 | 0 |
| 63 | PHE | 113 | ALA | -97.13 | 3.01 | 0 |
| 64 | ASP | 160 | PHE | -98.61 | 1.84 | 0 |
| 70 | ALA | 77 | VAL | -108.43 | 1.58 | 0 |
| 73 | LYS | 76 | GLY | -105.58 | 5.98 | 0 |
| 86 | SER | 214 | PRO | -60.33 | 4.58 | 0 |
| 87 | GLY | 216 | PHE | -112.12 | 2.4 | 0 |
| 88 | LEU | 93 | ALA | 100.99 | 2.21 | 0 |
| 89 | GLY | 92 | GLU | 105.76 | 2.08 | 0 |
| 105 | LEU | 174 | PRO | -94.13 | 5.57 | 0 |
| 107 | LYS | 175 | GLY | -97.44 | 7.11 | 0 |
| 108 | VAL | 174 | PRO | -85.78 | 4.17 | 0 |
| 109 | ALA | 112 | ALA | 118.54 | 3.92 | 0 |
| 109 | ALA | 172 | PRO | -69.28 | 4.01 | 0 |
| 120 | LEU | 125 | ALA | 112.71 | 4.92 | 0 |
| 130 | LYS | 139 | PHE | 90.23 | 3.66 | 0 |
| 131 | GLU | 137 | PHE | -116.86 | 2.55 | 0 |
| 132 | ALA | 139 | PHE | 124.2 | 5.16 | 0 |
| 139 | PHE | 160 | PHE | -71.54 | 4.55 | 0 |
| 166 | GLU | 169 | PHE | 97.75 | 4.71 | 0 |
| 173 | GLY | 203 | GLU | -86.78 | 1.91 | 0 |
| 175 | GLY | 200 | ARG | 116.01 | 3.67 | 0 |
| 188 | LYS | 191 | GLY | 124.28 | 4.26 | 0 |
| 242 | ARG | 245 | HIS | 69.75 | 4.21 | 0 |
| 255 | GLY | 258 | ARG | 119.54 | 2.85 | 0 |

*The value of energy should be less than 2.2 and Chi3 should be in between −87 and +97 degree

**Supplementary Table 10**: Energy scores of the top 10 vaccine-receptor docked complexes

| **Cluster** | **Members** | **Representative** | **Weighted Score** |
| --- | --- | --- | --- |
| 0 | 58 | Center | -1233.2 |
|  | 58 | Lowest Energy | -1445.5 |
| 1 | 39 | Center | -1211.9 |
|  | 39 | Lowest Energy | -1432.5 |
| 2 | 33 | Center | -1249.7 |
|  | 33 | Lowest Energy | -1368.2 |
| 3 | 32 | Center | -1580.3 |
|  | 32 | Lowest Energy | -1602.7 |
| 4 | 27 | Center | -1181 |
|  | 27 | Lowest Energy | -1446.7 |
| 5 | 26 | Center | -1467.5 |
|  | 26 | Lowest Energy | -1626.2 |
| 6 | 24 | Center | -1164.3 |
|  | 24 | Lowest Energy | -1610.9 |
| 7 | 23 | Center | -1158.3 |
|  | 23 | Lowest Energy | -1314.1 |
| 8 | 22 | Center | -1223 |
|  | 22 | Lowest Energy | -1375.1 |
| 9 | 21 | Center | -1431 |
|  | 21 | Lowest Energy | -1520.5 |
| 10 | 21 | Center | -1286 |
|  | 21 | Lowest Energy | -1353.4 |
| 11 | 18 | Center | -1308.5 |
|  | 18 | Lowest Energy | -1458 |
| 12 | 18 | Center | -1207.9 |
|  | 18 | Lowest Energy | -1337.9 |
| 13 | 17 | Center | -1328.2 |
|  | 17 | Lowest Energy | -1328.2 |
| 14 | 16 | Center | -1209.7 |
|  | 16 | Lowest Energy | -1290.1 |
| 15 | 16 | Center | -1262 |
|  | 16 | Lowest Energy | -1302.4 |
| 16 | 15 | Center | -1254.6 |
|  | 15 | Lowest Energy | -1378.5 |
| 17 | 15 | Center | -1292.4 |
|  | 15 | Lowest Energy | -1361.9 |
| 18 | 14 | Center | -1199.2 |
|  | 14 | Lowest Energy | -1391.7 |
| 19 | 14 | Center | -1383.3 |
|  | 14 | Lowest Energy | -1383.3 |
| 20 | 13 | Center | -1186.1 |
|  | 13 | Lowest Energy | -1355.7 |
| 21 | 13 | Center | -1180.9 |
|  | 13 | Lowest Energy | -1257.2 |
| 22 | 12 | Center | -1211 |
|  | 12 | Lowest Energy | -1448 |
| 23 | 12 | Center | -1340.5 |
|  | 12 | Lowest Energy | -1356.4 |
| 24 | 11 | Center | -1218.6 |
|  | 11 | Lowest Energy | -1321.1 |
| 25 | 11 | Center | -1189.6 |
|  | 11 | Lowest Energy | -1472.7 |
| 26 | 11 | Center | -1244.3 |
|  | 11 | Lowest Energy | -1300.4 |
| 27 | 11 | Center | -1220.6 |
|  | 11 | Lowest Energy | -1307.6 |
| 28 | 10 | Center | -1268.1 |
|  | 10 | Lowest Energy | -1405.7 |
| 29 | 10 | Center | -1251.9 |
|  | 10 | Lowest Energy | -1259 |

**Supplementary Table 11**: Molecular dynamic simulations of the vaccine-receptor complex

| **Time** | **Energy (kJ/mol)** | **Bond** | **Coulomb** | **VdW** | **RMSD** | | |
| --- | --- | --- | --- | --- | --- | --- | --- |
|  |  |  |  |  | **CA** | **Backbone** | **Heavy Atoms** |
| 0 | -9415434.125 | 178081.372 | -11950871.3 | 1778425.7 | 0.463 | 0.516 | 0.583 |
| 100 | -7314232.965 | 669242.903 | -10156784.82 | 1333727.5 | 1.702 | 1.735 | 1.889 |
| 200 | -7269456.868 | 644410.403 | -10045161.11 | 1298549.2 | 1.962 | 1.994 | 2.164 |
| 300 | -7241968.93 | 632628.18 | -9988998.167 | 1287578.1 | 2.101 | 2.127 | 2.324 |
| 400 | -7229977.885 | 620683.874 | -9959653.715 | 1285052 | 2.072 | 2.103 | 2.322 |
| 500 | -7222629.553 | 621859.157 | -9940511.206 | 1275587.5 | 2.3 | 2.335 | 2.536 |
| 600 | -7219610.443 | 611683.705 | -9931495.633 | 1280475.1 | 2.232 | 2.264 | 2.485 |
| 700 | -7217485.764 | 613563.05 | -9926372.982 | 1275283.2 | 2.474 | 2.502 | 2.732 |
| 800 | -7214014.14 | 613597.245 | -9921668.908 | 1273428.4 | 2.588 | 2.61 | 2.848 |
| 900 | -7211371.894 | 617018.565 | -9920510.653 | 1272751.2 | 2.535 | 2.561 | 2.806 |
| 1000 | -7212017.694 | 616359.795 | -9917407.205 | 1271229.3 | 2.65 | 2.677 | 2.93 |
| 1100 | -7209527.03 | 616154.87 | -9919229.245 | 1273832 | 2.697 | 2.723 | 2.971 |
| 1200 | -7213079.898 | 619258.261 | -9913222.591 | 1262601.4 | 2.863 | 2.893 | 3.122 |
| 1300 | -7213455.273 | 612599.447 | -9912787.669 | 1269027.4 | 3.187 | 3.212 | 3.433 |
| 1400 | -7218247.114 | 612340.932 | -9917809.499 | 1268430.6 | 3.206 | 3.236 | 3.465 |
| 1500 | -7216739.446 | 623205.387 | -9919875.199 | 1263615 | 3.239 | 3.274 | 3.487 |
| 1600 | -7212776.331 | 610056.139 | -9916754.878 | 1276613.8 | 3.25 | 3.282 | 3.518 |
| 1700 | -7214955.511 | 610577.161 | -9917837.877 | 1274279.2 | 3.342 | 3.376 | 3.602 |
| 1800 | -7214495.746 | 612898.507 | -9923553.347 | 1278144.4 | 3.544 | 3.577 | 3.79 |
| 1900 | -7213734.524 | 611351.434 | -9920470.461 | 1276801 | 3.347 | 3.381 | 3.618 |
| 2000 | -7212093.087 | 615130.306 | -9909051.689 | 1267025.6 | 3.337 | 3.364 | 3.64 |
| 2100 | -7213497.217 | 612140.021 | -9916692.988 | 1273296.6 | 3.498 | 3.528 | 3.775 |
| 2200 | -7214475.245 | 609403.116 | -9912144.819 | 1269311.1 | 3.917 | 3.946 | 4.161 |
| 2300 | -7212815.924 | 616505.893 | -9906910.266 | 1262059.9 | 3.886 | 3.909 | 4.133 |
| 2400 | -7214879.568 | 615737.672 | -9915163.415 | 1268184.6 | 3.981 | 4.008 | 4.233 |
| 2500 | -7215951.938 | 616335.676 | -9919373.083 | 1269005.7 | 3.71 | 3.74 | 3.972 |
| 2600 | -7214500.028 | 619160.257 | -9923286.372 | 1271019.2 | 3.956 | 3.99 | 4.186 |
| 2700 | -7218997.812 | 614288.037 | -9927921.53 | 1276119.2 | 4.166 | 4.195 | 4.396 |
| 2800 | -7213749.604 | 607168.225 | -9917049.574 | 1278525.4 | 4.058 | 4.088 | 4.316 |
| 2900 | -7213973.105 | 614508.415 | -9923536.685 | 1278422.3 | 4.013 | 4.045 | 4.277 |
| 3000 | -7212556.046 | 616073.089 | -9913282.89 | 1268379.7 | 4.165 | 4.195 | 4.403 |
| 3100 | -7215449.344 | 612404.96 | -9918902.636 | 1274059.5 | 3.864 | 3.898 | 4.101 |
| 3200 | -7215261.513 | 611999.066 | -9920759.119 | 1276253.7 | 3.863 | 3.893 | 4.102 |
| 3300 | -7210466.197 | 614420.831 | -9915316.927 | 1273294.3 | 3.87 | 3.899 | 4.138 |
| 3400 | -7212116.306 | 618230.206 | -9913816.786 | 1267072.5 | 3.956 | 3.99 | 4.199 |
| 3500 | -7212535.712 | 608341.008 | -9913428.474 | 1276485.1 | 4.147 | 4.178 | 4.385 |
| 3600 | -7216991.691 | 614057.808 | -9920154.051 | 1271515 | 4.276 | 4.307 | 4.506 |
| 3700 | -7214005.755 | 611706.378 | -9912727.916 | 1272045.5 | 4.252 | 4.285 | 4.487 |
| 3800 | -7218233.209 | 611311.193 | -9920083.263 | 1274921.7 | 4.201 | 4.231 | 4.453 |
| 3900 | -7213110.068 | 613737.699 | -9916699.559 | 1272270 | 4.552 | 4.581 | 4.766 |
| 4000 | -7214281.855 | 613407.315 | -9917175.133 | 1273297.2 | 4.629 | 4.655 | 4.846 |
| 4100 | -7214986.205 | 610255.119 | -9917064.814 | 1274101.4 | 4.361 | 4.387 | 4.585 |
| 4200 | -7215765.504 | 616684.222 | -9913760.089 | 1263113.8 | 4.447 | 4.476 | 4.671 |
| 4300 | -7212149.225 | 613874.3 | -9918118.204 | 1272658.9 | 4.105 | 4.132 | 4.347 |
| 4400 | -7216139.371 | 615274.79 | -9924605.31 | 1274032.4 | 4.235 | 4.263 | 4.472 |
| 4500 | -7216682.012 | 616337.24 | -9918210.901 | 1269072.9 | 4.464 | 4.485 | 4.695 |
| 4600 | -7218033.334 | 608883.84 | -9924497.784 | 1280469.1 | 4.349 | 4.379 | 4.587 |
| 4700 | -7218595.142 | 615475.9 | -9920802.785 | 1270425.4 | 4.837 | 4.566 | 5.047 |
| 4800 | -7216514.181 | 608451.212 | -9923045.388 | 1282577.9 | 4.549 | 4.374 | 4.798 |
| 4900 | -7216212.818 | 613801.681 | -9920340.819 | 1272240.1 | 4.719 | 4.445 | 4.947 |
| 5000 | -7215264.999 | 613216.3 | -9918443.593 | 1273442.7 | 4.906 | 4.235 | 5.131 |
| Mean | -7261872.454 | 607566.513 | -9968104.77 | 1284708.4 | 3.550 | 3.549 | 3.792 |
| Min | -9415434.125 | 178081.372 | -11950871.3 | 1262059.9 | 0.463 | 0.516 | 0.583 |
| Max | -7209527.03 | 669242.903 | -9906910.266 | 1778425.7 | 4.906 | 4.935 | 5.131 |
